# Structural Basis of Cold and Menthol Sensing by TRPM8

**DOI:** 10.1101/2025.09.09.675254

**Authors:** Hyuk-Joon Lee, Cheon-Gyu Park, Justin Gerald Fedor, Wyatt A. Peele, Mario J. Borgnia, Seok-Yong Lee

## Abstract

The transient receptor potential melastatin member 8 (TRPM8) is a polymodal ion channel that senses cold and menthol in mammals. Despite prior structural studies, the mechanisms by which cold and menthol activate TRPM8 remain unresolved. Here, we present cryo-EM structures representing the cold and menthol-dependent activation trajectories, combined with extensive functional analyses. We captured snapshots of cooling-dependent pore opening, which involves dramatic pore rearrangement, suggesting a mechanism for cold sensing. Moreover, menthol binds dynamically to induce channel activation, which may underlie menthol specificity for TRPM8. Finally, we show how TRPM8 integrates multiple modalities (cold and menthol) through overlapping but non-identical pathways, revealing the temperature-specific “cold spot”. These findings enhance our understanding of the molecular basis of physically and chemically induced cool sensation in mammals.

## Introduction

TRPM8 is the principal molecular sensor for cold and menthol, as evidenced by knockout studies. ^1–5^. It is an emerging therapeutic target for conditions such as dry eye, cold pain, and cough, highlighted by the recent FDA approval of a TRPM8 agonist for dry eye disease ^6–12^. Unlike other classes of sensory receptors, TRPM8 – a thermosensitive TRP (thermoTRP) sensory receptor – integrates two distinct modalities: physical (cold) and chemical (menthol) stimuli. This polymodal sensing is central to shaping our somatosensation, yet how a single receptor enables such polymodal sensing, and integrates these signals at the molecular level, remains a central question in mammalian thermosensation. In addition to cold and cooling agonists, TRPM8 requires PIP_2_ for its activation ^13^ and voltage enhances channel activity ^14^. Despite structural insights into ligand- and PIP_2_-dependent gating ^15–19^, major gaps remain. First, cooling-induced TRPM8 opening has not been captured and visualized. Second, while the polymodal gating of TRPM8 can be explained through allostery, the mechanistic link between cold and menthol sensation remains unclear ^14,20,21^. For instance, it is unknown for TRPM8 whether menthol and cold induce the same open state, utilize shared conformational pathways, and what their relative contributions are when co-applied. Third, there appears to be greater evolutionary pressure for TRPM8 to sense menthol than cold, as evidenced by the fact that TRPM8 from hibernating animals and mammoths has lost sensitivity to cold but retains sensitivity to menthol ^22–24^, in contrast to the species-dependent capsaicin sensitivity of TRPV1 ^25^. While TRPM4 responds to icilin, a large cooling agonist (M.W. 311 Da) ^26^, it remains unclear if menthol is specific to TRPM8 within the TRPM family and if so, how such a small molecule (M.W. 156 Da) can specifically trigger TRPM8 activation. Lastly, a recent structural study of avian and human TPRM8 proposes that menthol- and cold-induced channel opening involves an unusual “semi-swap” quaternary arrangement, especially for avian TRPM8 ^27^. This proposed cold and menthol open state supposedly adopts a gate distinct from that induced by other cooling agonists, but lacks functional validation. To address these questions, we determined cryo-electron microscopy (cryo-EM) structures of the *Mus musculus* TRPM8 channel (mTRPM8), representing the cold- and menthol-activated conformational landscapes, revealing cold-driven conformational changes at the gate and outer pore, as well as menthol-dependent conformational changes at the menthol binding pocket. Together with our extensive functional studies, we provide a mechanistic framework for how these two stimuli are integrated by TRPM8 to shape cooling sensation in mammals.

## Results

### Activation of TRPM8 by cold and menthol

Despite cold and menthol being capable of activating mammalian TRPM8 separately (Fig. 1A), validated open-channel structures by cold or menthol alone have been unattainable. As a polymodal channel, TRPM8 integrates multiple stimuli (*i.e.* cold, menthol, and membrane potential) in an allosteric manner to induce activation ^14,20^. For instance, when co-applied with cold, a sub-activating amount of menthol (1 µM) can further stimulate cold-induced channel currents (Fig. 1B,C) without affecting its temperature sensitivity Q_10_ (Fig. 1C). Increasing amounts of co-applied menthol (10 µM and 200 µM) at 8°C show currents saturate at ≤10 µM (Fig. 1D). Relative to a menthol EC_50_ of 60 µM at 20°C (fig. S1A-B), this saturating effect indicates a leftward shift in menthol affinity at 8°C by about 10-fold, consistent with a sensitizing effect between cold and menthol. We therefore sought to better understand this synergism and coupling to tease apart menthol- and cold-dependent transitions through extensive structural and biochemical studies.

**Figure 1.**
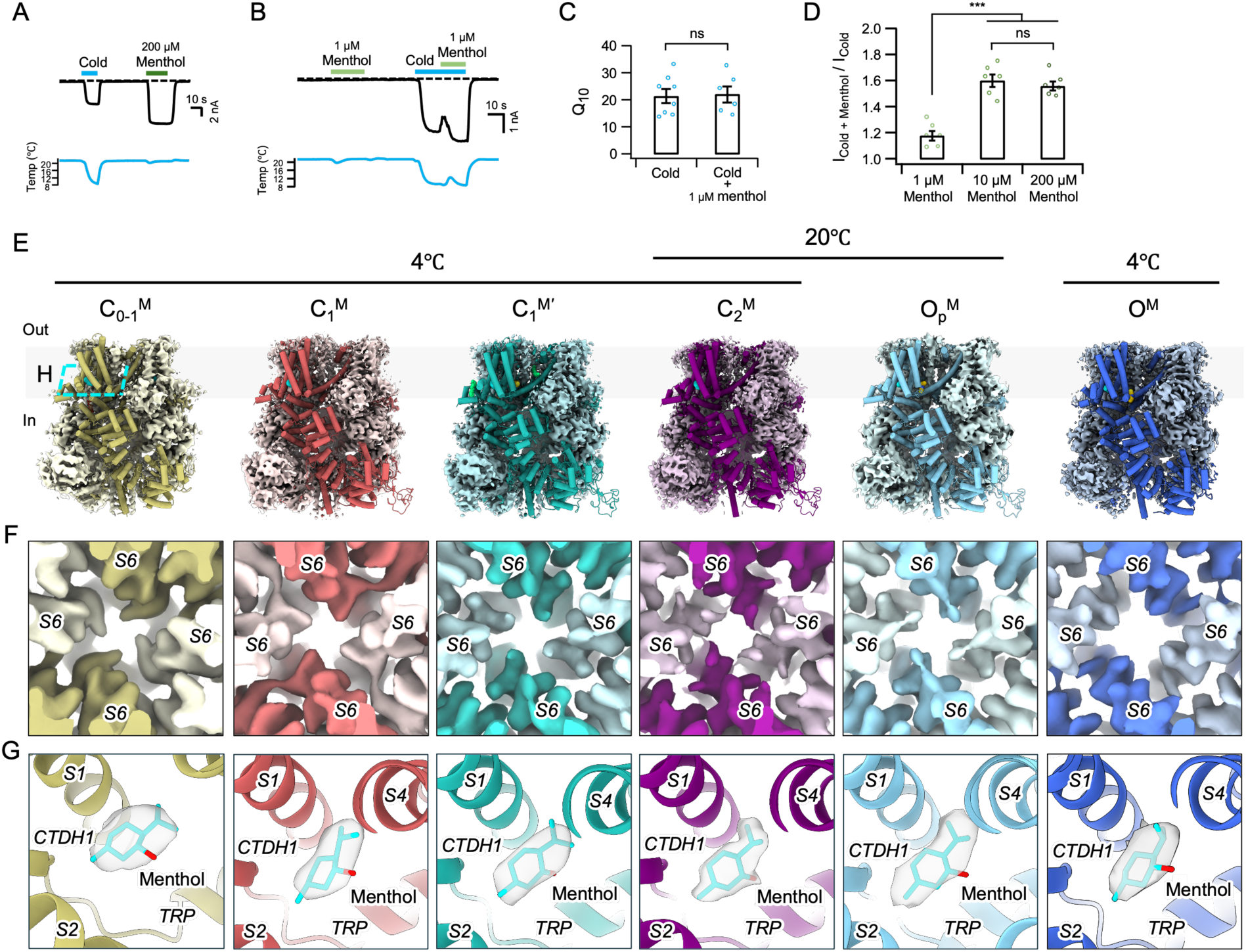
Overview of menthol and cold opening of TRPM8. **A)** Representative current traces of the mTRPM8 WT at −60 mV in HEK293T cells, elicited by cold alone or 200 µM menthol. **B)** Representative current traces, as in **A** demonstrating that a sub-activating concentration of menthol (1 µM) sensitizes mTRPM8 to cold. Horizontal colored lines represent the application of cold (sky blue), 1 μM menthol (light green), and 200 μM menthol (green), as indicated. Dotted lines indicate the zero-current level. **C)** Summary of the current ratios at −60 mV for mTRPM8 WT elicited by cold plus 1 µM, 10 µM and 200 μM menthol relative to cold alone. Currents were obtained following cold alone (*n* = 7 biological replicates) or cold plus menthol (*n* = 5 biological replicates). **D)** Q_10_ values for mTRPM8 WT activated by cold alone (*n* = 8 biological replicates) or in the presence of 1 μM menthol (*n* = 6 biological replicates) at -60 mV in HEK293T cells. For **C** and **D,** dots indicate the biological replicates for each experiment. \*\*\**P* < 0.001, student’s two-tailed unpaired *t*-test. Data are mean ± SEM. **E)** Overview of cryo-EM maps and 3D models for menthol-bound structures of mTRPM8 shown as sideview and colored according to the conformational assignment as indicated. Menthol is shown as cyan spheres in the boxed region. **F**) Bottom-up view of the EM density at the pore shows progressive dilation. **G**) View of the menthol binding site and menthol density for the respective states.

### Structural determination reveals multiple conformational states

TRPM8 is a homotetrameric channel composed of the transmembrane channel domain (TMD), which includes transmembrane helices S1 to S6, and the cytoplasmic melastatin homology regions (MHR) 1 to 4 (fig. S2A,B). Within the TMD, S1-S4 form a voltage-sensor-like domain (VSLD), while S5 and S6 form a central pore down the four-fold symmetry axis. The TMD of TRPM8 adopts a domain-swapped arrangement: the VSLD of each subunit interacts with S5 and S6 from the adjacent subunit’s (S5’ and S6’, fig. S2A,C). We purified mTRPM8 in glyco-diosgenin (GDN) detergent and determined structures of it in complex with menthol under various conditions (including at 20°C or 4°C). This allowed us to capture substates along the full breadth of the gating pathway (Fig. 1E), leading to the fully open state, as judged by the pore conformation (Fig. 1F). To describe the menthol- and cold-dependent conformational landscape of mTRPM8 we use a similar structural nomenclature as in our previous cryo-EM structures of mTRPM8 with the synthetic agonist Cryosim 3 (C3), allyl isothiocyanate (AITC), and PIP_2_ ^16^, determined at 20°C. These include three closed states: C_0_ (apo), C_1_ (PIP_2_ bound), C_2_ (PIP_2_ and C3), and an open state (O: C3, AITC and PIP_2_ bound), defined by distinct S6 gate residues ^16^.

Samples were prepared at 4°C unless otherwise described. We first incubated mTRPM8 with menthol and PIP_2_, which yielded a reconstruction in the C_1_ state (Fig. 1E and figs. S3–S5)^16^. We observed robust density corresponding to menthol, as with all our subsequent structures, and that PIP_2_ interacts with S4b, the TRP helix, and MHR4, as in the published PIP_2_-only C_1_ state determined at 20°C (Fig. 1E and fig. S6A,B). We therefore refer to this conformation as the C_1_^M^ state. However, capturing and visualizing the final open state of TRPM8 requires more than menthol, cold, and PIP_2_. Additional factors to induce activation are needed since TRPM8 exhibits very short open dwell times (< 1 ms) ^28^, neither menthol nor cold are strong agonists, it desensitizes, and cryo-EM conditions lack a transmembrane potential. We next prepared mTRPM8 with menthol, PIP_2_, and the activator rapamycin ^29^, density for which we did not observe. This yielded two 3D classes representing the C_1_^M^ state and one similar to the PIP_2_-free C_0_ state (Fig. 1E). Interestingly, in the latter state, PIP_2_ is bound but not as fully engaged as in C_1_, rather the inositol headgroup sits 4Å lower than in C_1_ and one acyl chain is inserted into the VSLD cavity (fig. S6A,B). This suggests a conformational state between the C_0_ and C_1_ states, which we term as C_0-1_^M^. Next, in addition to PIP_2_ and menthol – a type I cooling agonist – we included the type II cooling agonist, AITC, which binds at a separate site ^16,30^. This results in a conformation similar to the C_1_^M^, designated C_1_^M^’. We also determined the structure in complex with AITC and PIP_2_ only, which resulted in the C_1_ state, which we designated as C_1_^A^ (fig. S7A). We then introduced the Ile^846^Val mutation, which is known to increase menthol potency through enhanced interactions ^31^. Adding menthol and PIP_2_ to mTRPM8 Ile^846^Val resulted in a C_2_-like conformation, thus termed C_2_^M^. mTRPM8 Ile^846^Val was then treated at 20°C with AITC, menthol and PIP_2_, yielding two states: C_2_^M,20°C^ and another state with a dilated gate (Fig. 1F). As we will discuss later, this conformation is not in the open state, so we designate it as the pre-open state (O_P_^M^). Finally, we obtained the fully open state (O^M^) by preparing TRPM8 Ile^846^Val with AITC, menthol and PIP_2_ at 4°C, which also yielded another C_2_-like state, C_2_^M,4°C^. It is noteworthy that sub-activating concentration of AITC (0.5 mM) was used for all cryo-EM conditions, and we show in a later section (“AITC binds to the TRPM8 “coldspot” to mitigate channel desensitization”) that AITC has a substantial impact on cold activation.

Cursory inspection of the pore along these different states reveals a progressive dilation of the pore (Fig. 1F). All reconstructions are of excellent quality (3.1Å – 3.6 Å) (fig. S5, Table S1). Pore helix regions are less well resolved, and unsharpened cryo-EM maps are used to model this region (fig. S7C).

### Pore opening and S6 gate validation

We observe that the gradual rotation and structural changes of the pore-forming transmembrane helix (S6) from C_1_^M^ to O^M^ allows two narrow constriction points at Phe^979^ and Val^983^ to rotate away from the ion permeation pathway. This gradually dilates the S6 gate and introduces V976 as the gating residue (Fig 2A), consistent with the C3+AITC open structure resolved at 20°C previously ^16^. HOLE calculations indicate that the pore radius around Phe^979^ and Val^983^ expands in the O_P_^M^ state to slightly larger than 2 Å, then to ∼3 Å in the O^M^ state (Fig. 2B). We then functionally validated the S6 conformation and gate through cysteine scanning mutagenesis of S6 within the putative gate region (Fig. 2D–J, fig. S8). Here, different stimuli (menthol, C3, cold) induce currents in TRPM8 cysteine mutants before Cd^2+^ is added to the extracellular buffer. If the introduced cysteines face the ion conduction pathway, including a gate, they will react with Cd^2+^ and channel conductance will be affected. We previously utilized this approach to unambiguously determine the gate residue of TRPV4 ^32^. Consistent with our O^M^ structure, cold-induced currents for Val^972^Cys and Val^976^Cys were rapidly blocked by Cd^2+^, but not for WT, Leu^975^Cys nor Met^978^Cys (Fig. 2D,E). Positive voltages were used here because the conductance of these cysteine mutants is low due to their shift in voltage dependence (fig. S8A,B). Likewise, C3 and menthol activated mutants, Val^972^Cys and Val^976^Cys, were immediately blocked by Cd^2+^, while WT TRPM8, Leu^975^Cys and Met^978^Cys were not (Fig. 2F–I, fig. S8C). While we sought to cross-validate the open-state conformation by testing Phe^979^Cys, this mutant lacks channel activity. However, since Leu^975^ is one α-helical turn away from Phe^979^ in the O^M^ state, then lack of Cd^2+^ block in Leu^975^Cys unambiguously shows that Val^976^ is the gate residue in cold or cooling agonist open states, and that Phe^979^ is not. Interestingly, Leu^971^Cys, which is located toward the outer pore region, exhibits no block from cold or C3, but does show a slow block with menthol, indicating a difference in conformation or dynamics around this region when opened by menthol. Moreover, co-stimulated currents by cold and menthol are only partially blocked by Cd^2+^ (∼50%) (Fig. 2J). We therefore conclude that cold and cooling agonists induce a common gate architecture in TRPM8 consistent with our O^M^ structure, albeit with agonist-specific differences in the outer pore.

**Figure 2.**
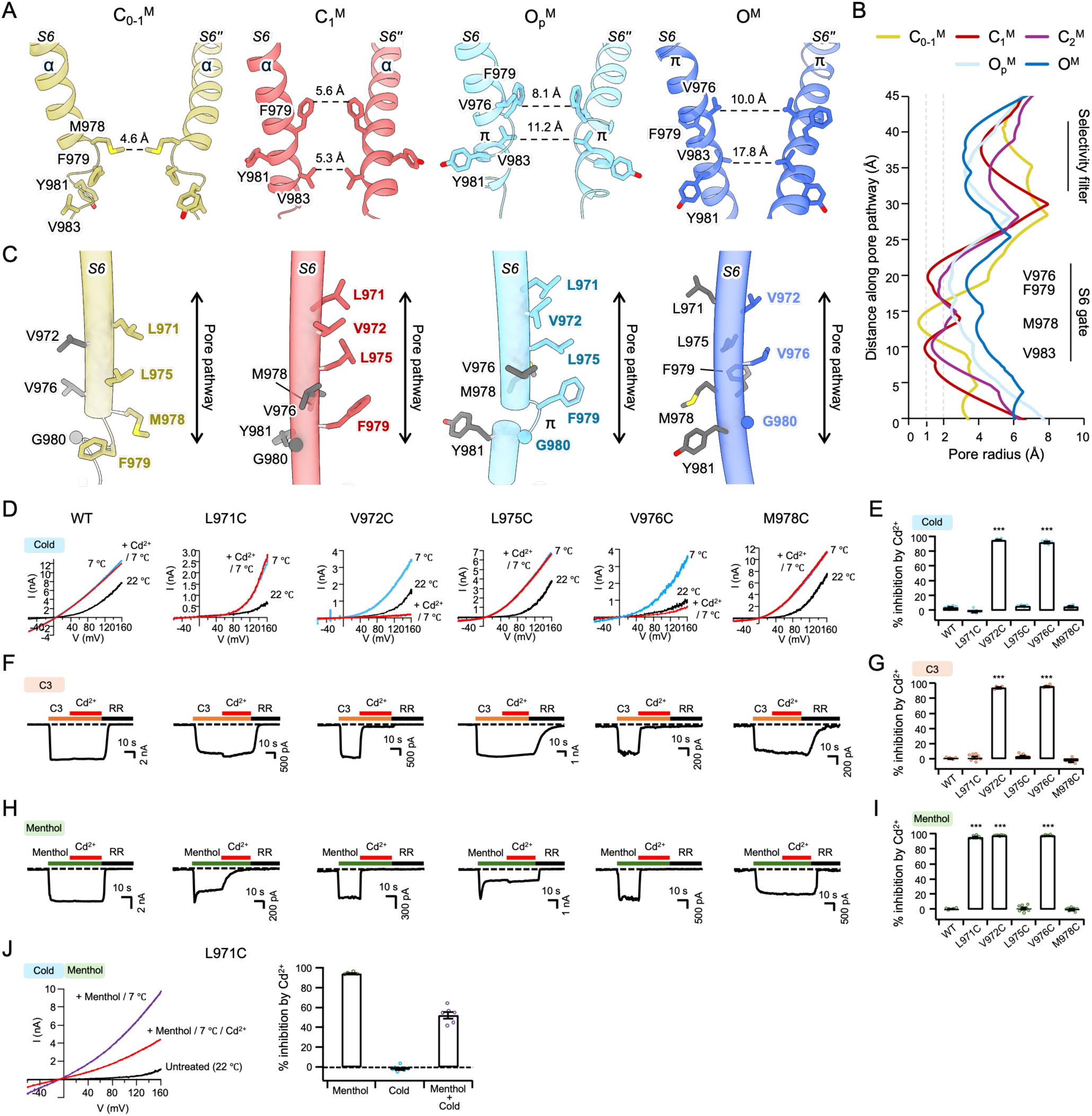
S6 gate structure and functional validation of the pore. **A)** Sideview of S6 helices from two opposite protomers (S6 and S6”) and conformation-specific gating residues for C_0-1_^M^, C_1_^M^, C_P_^M^, O^M^. **B)** HOLE analysis of pore diameter along pore length. Major constriction points as indicated for the selectivity filter and S6 gate. **C)** Progression from C_0-1_^M^ to O^M^ induces drastic conformational rearrangement of S6, which alters the pore lining residues with relative pore location indicated by the double arrow. Residues are shown as sticks and colored according to solvent accessibility (accessible – colored, buried – gray). **D)** Representative current-voltage (I–V) plots of the WT and cysteine mutant mTRPM8 channels in HEK293T cells, obtained using a 300 ms voltage ramp from -60 mV to +160mV, showing basal currents before any treatment (black trace), followed by initial application of cold stimuli (7°C) alone (blue trace), then co-application of cold stimuli with 20 μM Cd^2+^(red trace). **E)** Summary of inhibition of cold-evoked current by 20 μM Cd^2+^ measured at +160mV in HEK293T cell with WT and cysteine mutant mTRPM8 channels (*n* = 4–6 biological replicates). **F** and **H)** Representative currents of the WT and cysteine mutant mTRPM8 channels at -60 mV in HEK293T cells. Current traces elicited by 100 μM C3 (**F**, orange) or 200 μM menthol (**H**, green) and inhibited by subsequent application of 20 μM Cd^2+^ (red), then 50 μM ruthenium red (RR, black). Horizontal colored lines indicate the timing of compound application. Dotted lines indicate the zero-current level. **G** and **I)** Summary of % inhibition of C3-evoked (**G**) and menthol-evoked (**I**) current by 20 μM Cd^2+^ measured at -60mV in HEK293T cell with WT and cysteine mutant mTRPM8 channels (biological replicates, *n* = 3–7 for **G**, *n* = 5–7 for **I**). **J)** Representative current-voltage (I–V) plots of the mTRPM8 L971C cysteine mutant, as in **D,** showing basal currents before any treatment (black trace) at room temperature (22°C), followed by co-application of cold stimuli (7°C) and 200 μM menthol (blue trace), then finally co-application of cold stimuli, 200 μM menthol and 20 μM Cd^2+^(red trace). Summary of 20 μM Cd^2+^ inhibition on currents evoked by cold stimuli, 200 μM menthol, or their combination at +160mV in HEK293T cell with the mTRPM8 Leu^971^Cys cysteine mutant (*n* = 4-6 biological replicates). For bar charts in **E**, **G**, **I**, **J**, dots indicate the individual data points for each experiment. \*\*\**P* < 0.001, using one-way ANOVA followed by Dunnett’s post-hoc test. Data are mean ± SEM. Throughout the figures, amino acids are abbreviated: Ala (A), Cys (C), Asp (D), Glu (E), Phe (F), Gly (G), His (H), Ile (I), Lys (K), Leu (L), Met (M), Asn (N), Pro (P), Gln (Q), Arg (R), Ser (S), Thr (T), Val (V), Trp (W), Tyr (Y).

### Sequential conformational changes leading to cold- and menthol-induced opening

A comparison of the overall structures of the apo C_0_ (20°C, PDB 8E4P), and menthol-bound 4°C C_0-1_^M^, C_1_^M^, and C_2_^M^ states provides an overall conformational trajectory for mTRPM8 channel opening by cold and menthol (Fig. 3A). Upon menthol and full PIP_2_ binding, formation of a 3_10_ helix in the S4-S5 linker is accompanied by a 20° upward rotation of the TRP helix with resultant shifts in S5 and S6, leading to the C_1_^M^ then C_2_^M^ states. Thereafter, AITC binding facilitated full conversion of S6 to yield the O^M^ state at 4 °C. But with so many factors at play, it is unclear which of these conformational transitions can be attributable to cold versus ligand binding.

**Figure 3.**
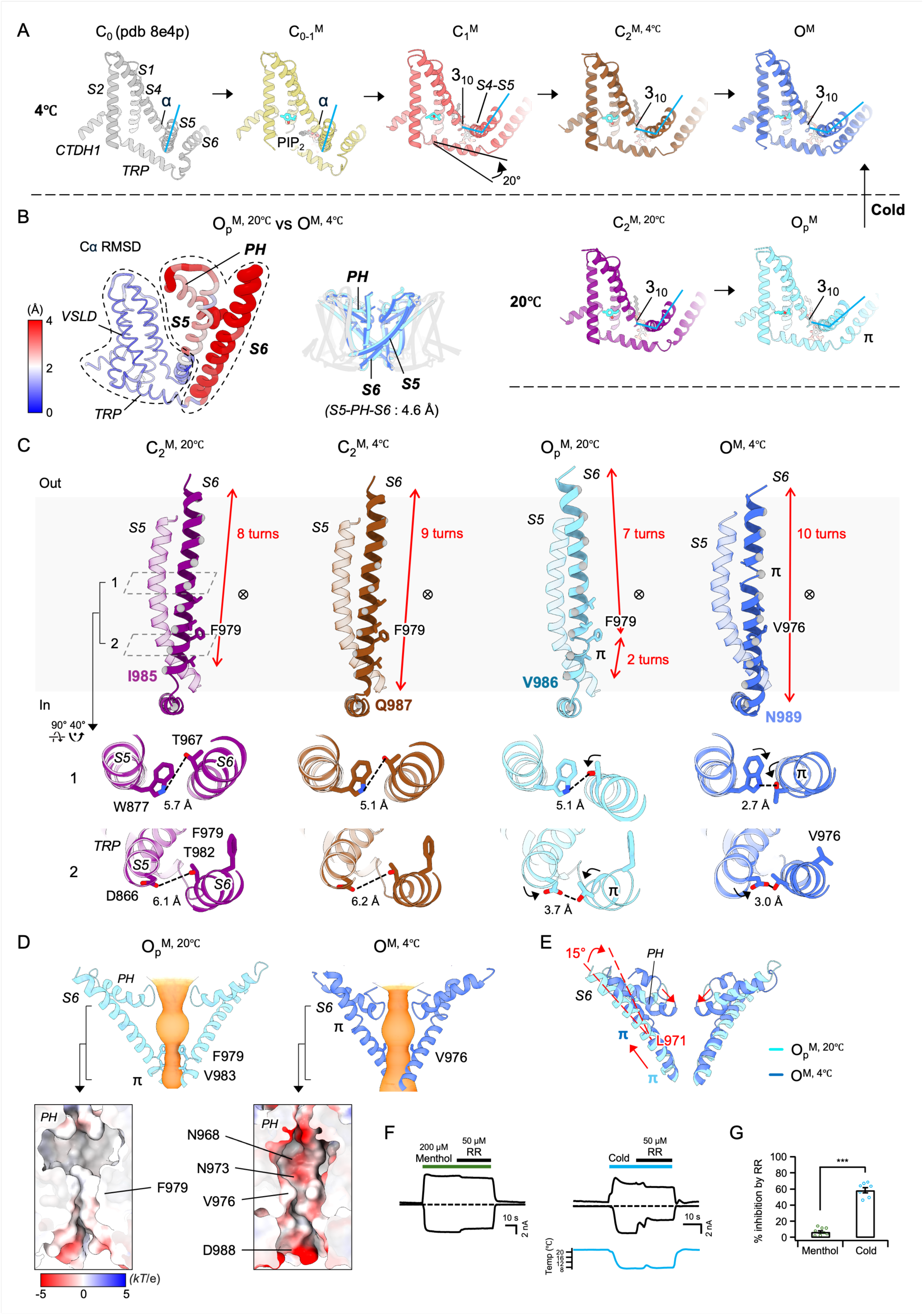
Cold-induced conformational changes in TRPM8 activation. **A)** Conformational trajectory of TRPM8 opening, represented by C_0_ (PDB 8E4P), C_0-1_^M^, C_1_^M^, C_2_^M,4°C^, and O^M^ at 4°C (top row), and comparison with C_2_^M,20°C^ and O_P_^M^ at 20°C (bottom row). Hallmark conformational changes are highlighted: binding of PIP_2_ and menthol, movement of the TRP helix, S4-S5/S5, and helical transition in S4B. **B)** The largest temperature-dependent conformational changes occur between O_P_^M^ and O^M^ and are limited to S6, PH and S5, as indicated by the cartoon thickness and color to denote Cα RMSD. **C)** Comparison of TRP-S6 and S5 between C_2_^M,4°C^ and C_2_^M,20°C^ then O_P_^M^ and O^M^ states. Gray spheres every four Cα help visualize conformational changes to S6. S6 lengthening is denoted by the red line with the terminal S6 residue labeled in bold, the respective gating residues are labeled along with pore-lining residues shown as sticks. The locations of the pore (crossed-circle mark) and π-bulge are indicated. Clipped extracellular views 1 and 2 show changes in hydrogen-bonding networks during cooling. Dashed lines indicate the distances between residues involved in those networks. **D)** The difference in pore helix conformation and pore cavity electrostatics (insets) between O_P_^M^ and O^M^. **E)** Upon cooling, O_P_^M^ converts to O^M^ through lengthening and rotation of S6, inducing movement of the π-bulge and PH. **F)** Representative currents of the WT mTRPM8 in HEK293T cells at -60 mV (lower traces) and +60 mV (upper traces) in response of currents induced by 200 µM menthol (left, green) or 7°C cooling (right, blue) and then 50 μM RR (black) as indicated. Dotted lines indicate the zero-current level. **G)** Summary of % inhibition of menthol- or cold-evoked inward currents (-60 mV) where data are mean ± SEM (*n* ≥ 7 biological replicates). Dots indicate the individual data points for each experiment with mean ± SEM. \*\*\**P* < 0.001, using two-way ANOVA followed by Sidak post-hoc test.

### Cold-induced transitions of S6 and the PH drive the final stages of TRPM8 activation

As previously shown ^16^, PIP_2_, along with menthol, plays an important role in driving the initial conformational changes to C_1_^M^, and introduction of the menthol potency-enhancing Ile^846^Val mutation facilitated transition to the C_2_^M^ state. In comparison, mTRPM8 with AITC and PIP_2_ or menthol and AITC remains in the C_1_ state, supporting the role of enhanced menthol binding to reach the C_2_^M^ state (fig. S6A). Therefore, we can attribute much of the early stages of the conformational pathway of TRPM8 to cumulative effects of menthol/PIP_2_. We therefore compared the structures obtained at 20°C and 4°C from the final set of biochemical conditions (mTRPM8 Ile^846^Val +menthol, PIP_2_, AITC), which yielded C_2_^M,20°C^, C_2_^M,4°C^, O_P_^M^ and O^M^ states (Fig. 3A). Strikingly, temperature dependent conformational changes are prominent between the O_P_^M^ (20°C) and O^M^ (4°C) states; the pore (S5, S6 and the PH) exhibit dramatic differences (overall Cα RMSD ∼4.6A), while there were no changes in the VSLD nor menthol pose (Fig. 3B,C). Transition from O_P_^M^ (20°C) to O^M^ (4°C) includes a substantial reorganizing of the TRP-S6 linker and lengthening of the S6 helix by one turn (Fig. 3C). Transition from C_2_^M,20°C^ to C_2_^M,4°C^ only adds half a helical turn to S6 due to TRP-S6 linker re-organization (Fig. 3C). Notably, conversion of C_2_^M,20°C^ to O_P_^M^ is marked by the formation of a π-bulge in S6 (M979-T983). During the cold-induced transition of O_P_^M^ to O^M^ this π-bulge migrates up S6 (I962-S966). Additionally, two H-bonds are formed in O^M^ that do not occur in C_2_^M^: Trp^877^(S5)–Thr^967^(S6) and Asp^866^(S5)–Thr^967^(S6) (Fig. 3C). The overall effect of these transitions is the extension, rotation and stabilization of S6 to open the pore.

An additional consequence of the movements of S6 are the associated motions of the PH (Fig. 3D, fig. S6C). Relative to O_P_^M^, the PH in O^M^ moves inward and down as S6 tilts by 15°, pivoting near Leu^971^ (Fig. 3E), increasing the electronegativity of the pore (Fig. 3D). These temperature-dependent conformational and electrostatic changes in the pore cavity are in line with the agonist-dependent effects on Cd^2+^ block of Leu^971^Cys (Fig. 2J). To additionally probe the outer pore structure, we co-applied 50 µM ruthenium red (RR) with menthol or cold stimulation of mTRPM8 in electrophysiology experiments (Fig. 3F). RR acts as a pore blocker for many TRP channels by interacting with the selectivity filter ^33,34^. Interestingly, we observed menthol-induced currents were largely unblocked, but cold-induced inward currents were most sensitive to RR block (Fig. F,G). These data are consistent with agonist-specific open-state conformations of the outer pore region (near the pore entry and the selectivity filter).

Our structure-based observation of cold affecting the later stages of the landscape to drive pore opening is in line with single-channel studies ^28^. Importantly, these cold-induced conformational changes in S6 and the PH dramatically change the pore conformation and its electronegativity.

### Menthol binds specifically to the VSLD cavity of TRPM8

Menthol binds within the VSLD cavity – located between the VSLD and the TRP helix (Fig. 4A), for which all our maps exhibit robust cryo-EM density and is absent in the published apo data (Fig. 1E, fig. S6C) ^16^. The C_2_^M^ dataset provides particularly excellent EM density, allowing unambiguous assignment of the menthol pose (Fig. 4A). Inside this cavity, menthol is surrounded by residues F1013 from the C-terminal domain helix 1 (CTDH1), Asn^741^, Val^742^, and Tyr^745^ from S1; Leu^778^ and Asp^781^ from S2; Arg^842^ and Ile^846^ from S4, and Tyr^1005^ and Arg^1008^ from the TRP helix (Fig. 4A inset). No residues from S3 participate in menthol binding. Functional analyses agree with prior studies on the roles of Tyr^745^, Arg^742^, Ile^846^ and the TRP helix in menthol binding, thus validating this as the menthol binding site (fig. S1D–E) ^21,31,35^.

**Figure 4.**
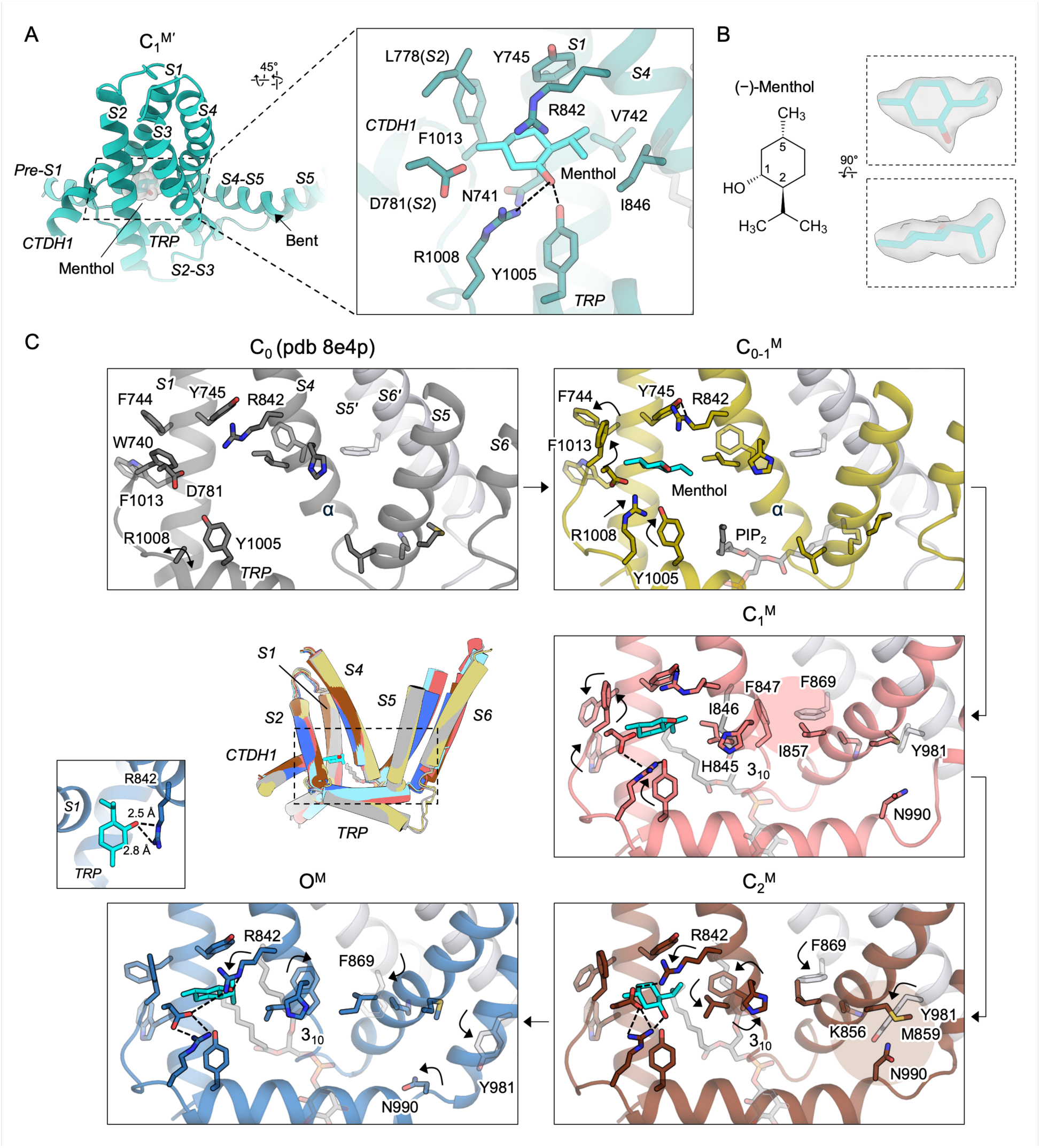
Menthol binding site and conformational changes leading to pore opening at 4°C. **A)** Menthol binds to the VSLD cavity flanked by the TRP helix. Interacting residues are shown as sticks and hydrogen bonds as dashed lines. **B)** Chemical structure for the (–)-menthol, its natural and active stereoisomer, and the unambiguous modeled pose for menthol (cyan sticks) in the C_1-2_^M^ EM density. **C)** Menthol binding pocket rearrangements at 4°C, as contextualized in the inset. Each pane depicts the progressive conformational changes initiating at the menthol site and leading to the S6 pore helix. Sidechains are shown as sticks and their movements indicated by black arrows. Putative hydrogen bonds are shown as black dashed lines. Conversion of a portion of the S4b helix from α to 3_10_ in C_1_^M^, C_2_^M^ and O^M^ is indicated. The propagation of sidechain interaction cluster rearrangements are highlighted in C_1_^M^ and C_2_^M^.

Other Type I cooling agonists, such as the menthol derivative WS-12, cryosim-3, and icilin, also bind within the VSLD cavity, but menthol sits deeper into the VSLD cavity, closer to CTDH1 and S1, and this unique positioning distinguishes it from those of other agonists (fig. S9A,B) ^9,16,36^. The menthol binding site of TRPM8 is thought to be conserved within the TRPM family (fig. S1F,G)^26^. However, we found that menthol alone does not activate TRPM2, TRPM3, or TRPM4, nor does it affect TRPM4 Ca^2+^ gating. (fig. S1H–K). This suggests that, within the TRPM family, menthol acts specifically on TRPM8, although menthol can activate TRPA1 ^37^. Of the residues lining the menthol site, Asp^781^ and Phe^1013^ are unique to mTRPM8 (Fig. 4A, fig. S1F). While Phe^1013^Ala substantially reduces menthol-induced currents in mTRPM8 (fig. S1E), the Glu^827^Asp/Pro^1077^Phe mutations in hTRPM4 do not confer menthol sensitivity (fig. S1I,J). This suggests specificity to menthol in TRPM8 is not simply conferred by sequence conservation or a particular cavity architecture but may instead require accommodation of menthol and/or cavity dynamics to induce long-range conformational changes via a specific network of interactions.

### Dynamic binding of menthol helps drive early-stage conformational changes

With the latter conformational stages being the most temperature sensitive, we then inspected the early stages to more deeply probe menthol binding and the conformational changes that lead to pore opening. We therefore compared the state-dependent conformational changes among our 4°C structures by alignment of their VSLDs: C_0-1_^M^, C_1_^M^, and C_2_^M,4C^ (Fig. 4C). We first note that in the apo C_0_ state at 20°C (PDB 8E4P), F1013 (CTDH1), F744 (S1), and W740 (S1) form an aromatic cluster, which rearranges upon menthol binding (Fig. 4C). Meanwhile, TRP helix residues Tyr^1005^ and Arg^1008^ move to engage menthol in the C_0-1_^M^ state, which is followed by a 20° upward swing of the TRP helix and an α-to-3_10_ helical transition at S4b in the C_1_^M^ state (Fig. 4C). Subsequent rotation of S4b establishes the Arg^842^-Asp^781^-Arg^1008^ triad, which remains stable from the C_2_^M^ state through to O^M^ and likely enhances the coupling between S4 and the TRP helix. PIP_2_ binding also contributes to the structural transitions from C_0_ to C_1_ ^13,16,18^. Notably, menthol itself shifts and substantially rotates (∼45 °) during the C_0-1_^M^ to C_1_^M^ transition, followed by a ∼-50° rotation upon transition to C_2_^M^, then another ∼50° rotation to the O_P_^M^/O^M^ states. Formation of a hydrogen bond with Arg^842^ in the O_P_^M^/O^M^ states suggests increasing menthol binding affinity (Fig. 4C inset). This, and the ability of menthol to conformationally adapt to the induced protein structural changes, is consistent with an induced fit binding mechanism.

Overall, menthol-induces sequential rearrangements that propagate from CTDH1/S1 through the TRP helix to S4, as facilitated by PIP_2_ (fig. S10). This signal is propagated further when rotation of S4b disrupts a critical interaction network formed by Phe^847^ (S4b), Ile^857^ (S4-S5), and Phe^869^ (neighboring S5; S5’) in the C_1_^M^ state. A new interaction cluster involving Lys^856^ (S4-S5), Asn^990^ (TRP helix), and Tyr^981^ (S6’) then stabilizes the C_2_^M^ state (Fig. 4C). This cluster then weakens in O^M^, where disengagement of Tyr^981^ (S6’) and Asn^990^ (TRP) facilitates rotation of S6’ (Fig. 4C).

### Distinct but overlapping networks for menthol and cold activation

We performed electrophysiology experiments on alanine mutants of conformationally variable residues in the VSLD, TRP helix, S4-S5 linker, S5, and S6. Menthol- or cold-induced inward current densities were normalized to WT and plotted (Fig. 5A, fig. S11A). Mutants appearing above or below the line of identity more substantially affect cold- or menthol-induced currents, respectively, thus termed “cold preferred” or “menthol preferred”. Mutants with ≥70% WT activity for both menthol and cold are “non-affectors”. Mutants appearing at the origin are “critical (non-specific)” (Fig. 5A). Plotting the WT-normalized inward current ratio [(I_cold_/I_menthol_)/(I^WT^ /I^WT^)] recapitulates the categorization and gives fold-difference of cold-to-menthol preference compared to WT, where non-effectors are found ±25% of WT (Fig. 5B). Of the critical (non-specific) mutants, five had measurable outward current (fig. S11B,C), and the remainder were confirmed to be surface expressed but functionally dead (fig. S11D). We first mapped the menthol preferred and critical (non-specific) residues onto the O^M^ structure (Fig. 5C). Many functionally important residues form a cluster around the menthol binding site to S4 within the VSLD and another around the S4-S5 linker and S6. Mapping of the cold preferred and critical (non-specific) residues reveals a continuous interaction network extending from within the VSLD to S6, including a distinct subset of residues at S4, the S4-S5 linker, S5’, and S6’, illuminating how cold sensing is transmitted to the gate. Notably, the cold-preferred network involves Met^859^, Gln^861^, and Arg^862^ from the S4-S5 linker, as well as Tyr^981^ from S6’. These are a separate set of residues from the “menthol preferred” residues Leu^860^ (S4-S5 linker), Phe^870^ (S5’), and Leu^970^ (S6’). These data suggest that, although cold- or menthol-induced signals propagate through shared structural elements, they do so via largely different sets of residues that interact via different networks to open the pore.

**Figure 5.**
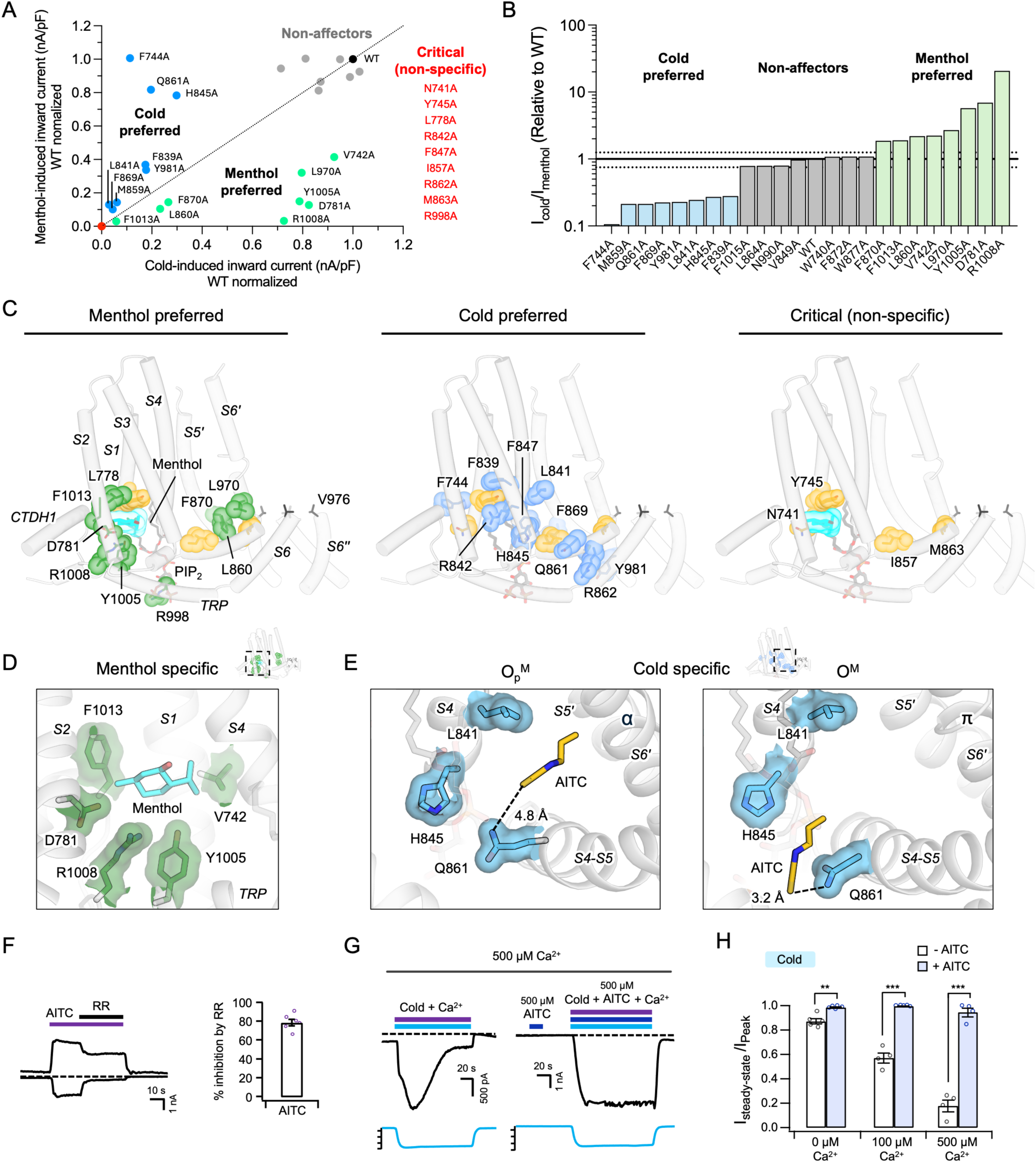
Functional mapping of menthol- and cold-conformational networks. **A)** Plot of mTRPM8 alanine mutants as characterized by whole-cell electrophysiology (-60 mV, inward current density) where cold-activated (≤10°C, x-axis) and menthol-activated (200 µM, y-axis) responses are normalized to those of WT mTRPM8. Residues plotted above the line of identity more greatly affect cold sensing (cold preferred) and those below the line of identity more greatly affect menthol sensing (menthol preferred). Several non-functional mutants have no measurable inward currents, as indicated. **B)** Plot of inward current density ratio for cold:menthol activation for mTRPM8 alanine mutants set relative to WT. An arbitrary threshold of ±25% from WT indicates residues not substantially impacted by mutation to alanine (non-affectors). Mutants below or above these thresholds are cold or menthol dominant, respectively. **C)** Mapping of functionally important residues onto an O^M^ protomer TMD, as determined from the above plots. Menthol is shown in cyan sticks and spheres, PIP_2_ as grey sticks, and pore residue Val^976^ as black sticks. Menthol or cold preferred residues are shown as green or blue spheres and sticks, respectively. Residues that show no inward or outward currents are mapped as “Critical (non-specific)”. **D)** Menthol binding site with menthol in cyan sticks and menthol preferred residues in green surface and sticks. **E)** “Coldspot” residues shown as blue surface and sticks with AITC shown as yellow sticks in the O_P_^M^ and O^M^ structures. **F)** Right, representative currents of the WT mTRPM8 channels in HEK293T cells at -60 mV (lower traces) and +60 mV (upper traces) in response to 2 mM AITC (dark blue) and 50 μM RR (black) as indicated. Dotted lines indicate the zero-current level. Left, summary of % inhibition of AITC-evoked inward currents (-60 mV) where data are mean ± SEM (*n* = 6 biological replicates). **G)** Representative currents of WT mTRPM8 at –60 mV in HEK293T cells elicited by cold alone (sky blue) or with sub-activating concentrations of AITC (500 μM) (dark blue). Dotted lines indicate the zero-current level. **H)** Summary of remaining current after desensitization in the presence of 0 μM, 100 μM, and 500 μM extracellular Ca^2+^ (*n* = 4–5 biological replicates). Dots indicate the individual data points for each experiment with mean ± SEM. \*\**P* < 0.01, \*\*\**P* < 0.001, using two-way ANOVA followed by Sidak post-hoc test.

Careful inspection of Fig. 5A reveals residues that are highly specific in their effects to cold or menthol sensing, found in the upper left and lower right of the plot. Not surprisingly, the menthol specific residues line the menthol cavity (Fig. 5D). Interestingly, of the three highly cold specific mutants (Phe^744^Ala, Gln^861^Ala and His^845^Ala), Gln^861^ (S4-S5 linker) and His^845^ (S4) form a cluster with the cold preferred residue Leu^841^ (S4) adjacent the S6 π-bulge position in the cold-induced O^M^ state (Fig. 5E). This also happens to be where AITC binds, consistent with Gln^861^ being critical for AITC binding ^16^. Comparing the O_P_^M^ and O^M^ states, AITC itself appears to undergo a temperature-dependent shift in binding position (Fig. 5E). So despite cold-preferred residues being distributed throughout TRPM8, where S6 and the PH appear particularly temperature sensitive (Fig. 3B), ^24^ we identify this as a cold sensory hotspot, or “coldspot”, of TRPM8.

### AITC binds to the TRPM8 “coldspot” to mitigate channel desensitization

In a classical two-state model, cold temperatures are known to destabilize the closed state ^14^. Similarly, AITC - a type II cooling agonist - also destabilizes the closed state, while type I agonists such as menthol and C3 stabilize the open state ^16,30^. Therefore, AITC and cold exert similar effects on activation kinetics. We therefore hypothesized that AITC may stabilize conformational changes specific to cold activation, and that cold and AITC may at least partially utilize a shared activation mechanism. In accordance with AITC binding at the coldspot to stabilize a cold-specific conformation, RR partially blocks AITC-activated currents (Fig. 5F) similar with cold activation (Fig. 3F,G). Moreover, Cd^2+^ block of AITC-activated TRPM8 is consistent with Val^976^Cys and Val^972^Cys being pore-lining residues, as with activation by menthol, cold, voltage and C3 (fig. S8E,F). Next, since conformational rearrangement of the interface between the S4-S5 linker and S5’/S6’ is critical for desensitization ^15^ – which occurs rapidly (τ ≈ 3 s) in the presence of Ca^2+^ or AMG2850 (fig. S12A–D) – then AITC binding to this interface may help destabilize the desensitized state. Indeed, the addition of sub-activating concentrations of AITC (500 μM), does not induce opening but substantially reduces desensitization of both cold- and cooling agonist-elicited currents (Fig. 5G,H, fig. S12E–G). Notably, AITC has a much greater effect on cold desensitization such that cold-activated TRPM8 does not desensitize under increasing Ca^2+^ concentrations (Fig. 5H, fig. S12E). Therefore, AITC and cold share a common activation network, unique from menthol, that AITC acts upon by binding to the TRPM8 coldspot. More importantly, the use of AITC as a functional probe highlights the core role of this cold sensory network in channel activation and adaptation to stimuli.

## Discussion

Here, we delineated the cold- and menthol-dependent conformational landscape of mTRPM8. Our structural analyses suggest that menthol and PIP_2_ binding influence early conformational transitions, while cold affects later stages of the landscape to drive pore opening. Our cold-dependent conformational changes involve dramatic pore rearrangement, including extension and stabilization of S6. This aligns with previous electrophysiological characterization of TRPM8 wherein cold activation is accompanied by large negative enthalpy and negative entropy changes, indicating a transition toward a more stable and ordered open state ^38^. The coil-to-helix transition in S6 and formation of S6-stabilizing H-bonds suggests that thermally induced helix formation at lower temperatures contributes to cold sensing (Fig. 3 and 6) ^39^, thus providing a structural basis for cold sensing by TRPM8. Energetically, the formation of two helical turns plus two additional S6-stabilizing H-bonds per subunit (Fig. 3C), could provide ∼60–100 kcal/mol of stabilization through H-bonding, which would be amplified by the effect of cold temperatures on increasing H-bond strength ^40^. This is already comparable to the enthalpy associated with TRPM8 gating (–110 kcal/mol) ^20^, and restructuring of the pore architecture and electrostatics likely further contribute to the energetics of cold opening of TRPM8 ^41,42^.

**Figure 6.**
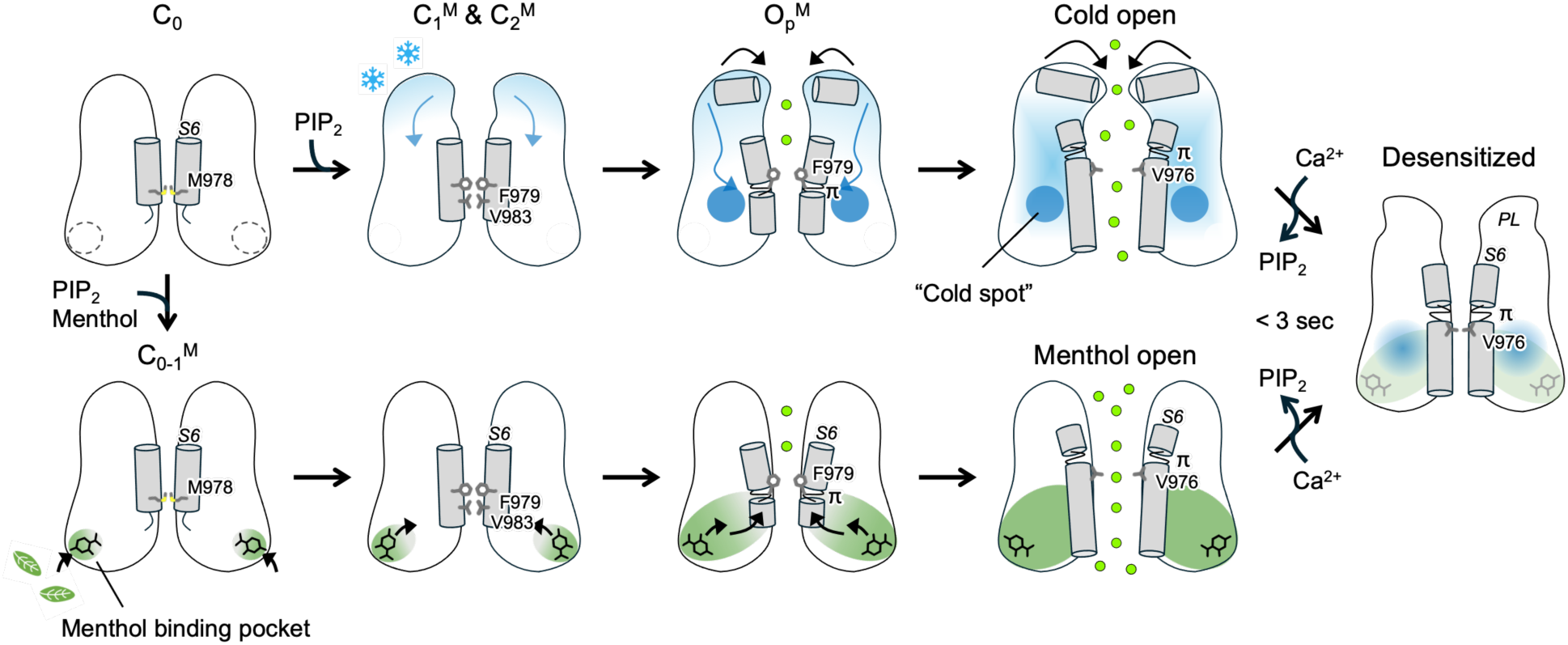
Model for cold and menthol activation of TRPM8. Starting from the apo state C_0_, cold temperatures activate in a delocalized fashion, originating from the outer pore region, propagating towards the “coldspot” to induce pore opening through rearrangements of S6 and the PH. Menthol activates TRPM8 by binding dynamically to the menthol binding site, initiating a sequence of conformational changes that propagate to S6 for gate opening via an overlapping but different set of residues compared to cold. The resulting open states share a common gate (Val^976^) but have unique outer pore structures. Prolonged stimulation of these open states causes rapid adaptation to a common desensitized state.

Our structure-guided mutagenesis and accessibility studies reveal that TRPM8’s polymodal gating arises from overlapping but distinct conformational pathways (Fig. 5C). Both physical (cold) and chemical (menthol) stimuli transmit forces through different networks of residues within the same structural modules (S4, the S4-S5 linker, S5, S6, and the TRP helix) to converge on the shared open S6 gate conformation. This explains their coupling and synergy in channel opening: if both stimuli utilize the same network or have distinct gate conformations, there will be no synergy. Despite this shared gate, cold and menthol induce different outer pore conformations. We also identified a cluster of residues that is specifically crucial for cold sensing, which we term the TRPM8 “coldspot”; to which AITC binds to stabilize the cold-specific conformational pathway, including greater stabilization away from cold desensitization. We therefore suggest that the cold sensory network converges onto the coldspot, situated at the S4-S5 linker and S5, because it is central to channel activation and stimulus adaptation (Fig. 6). That various stimuli activate TRPM8 through different interaction networks on shared structural elements raises the possibility of new or revised avenues for pharmacologically targeting TRPM8, through co-application of existing or engineered compounds to potentiate or bias signaling.

Unexpectedly, our structural analysis reveals gradual changes in the conformation of the menthol binding site, accompanied by adaptive menthol positioning that trigger a propagating conformational wave leading to pore opening (Fig. 6 and fig. S10). This induced-fit mechanism is distinct from the conformational selection mechanism employed by many ligand-gated ion channels ^43,44^. Dynamic menthol binding may favor higher-affinity conformational states until O_P_^M^, promoting the forward transition along the activation pathway ^45^. We propose that this dynamic induced-fit mechanism is evolutionarily conserved and underlies menthol specificity for TRPM8 within the TRPM family. Moreover, the strict requirement of PIP_2_ for TRPM8 activation likely contributes to the specificity for menthol by TRPM8 ^46^, as PIP_2_ is not a requirement for activation for other TRPM members, and its binding site may vary within the TRPM family ^47–50^.

Our findings contrast with a recent proposal that cold and menthol gating in TRPM8 involves a rearrangement between full-swap and semi-swap quaternary architectures, with the semi-swap conformation being essential for opening (fig. S2C and S13A)^27^. However, there are several issues with a semi-swap arrangement for open state TRPM8. First, all the TRPM8 structures in the semi-swapped arrangement, including the proposed cold/pH-open and menthol-bound states, exhibit a fully α-helical S6 with Phe^979^ as the gate residue (fig. S13B) ^27^. Importantly, we unequivocally demonstrate that Val^976^ is the gate residue for TRPM8 activated by cold, menthol, voltage and other cooling agonists, not Phe^979^ (Fig. 2D–J, fig. S13B). Only in this open state is the pore sufficiently dilated (>3 Å radius) to facilitate movement of hydrated cations (Fig. 2B). Second, for Val^976^ to become the gate residue, helical extension and rotation of S6 requires formation and movement of the π-bulge and tightening of the linkage between S6 and the TRP helix. This drastically contrasts with the loose linkage observed between the TRP helix and the short fully α-helical in the semi-swap arrangement, where the Phe^979^ residue gates a ∼ 2 Å radius pore (fig. S13B). Third, it was proposed that a semi-swap to full-swap transition is required for TRPM8 desensitization. Such a process involves large, concerted movements requiring at least partial tetramer disassembly, which occurs over minutes to hours according to the H–D exchange experiments (fig. S13C) ^27^. For instance, the tetramer-to-pentamer rearrangement of TRPV3 occurs with a time constant that exceeds 3 min ^51^. However, we demonstrate that TRPM8 desensitization from cold or menthol occurs within 3 s in the plasma membrane (fig. S12A–D). Additionally, we have previously demonstrated that TRPM8 inhibitors are conformationally selective, binding to a single desensitized state within the domain-swapped interface of S5’/S6’ and the S4-S5 linker and engaging the π-bulge of S6. All together it is highly unlikely TRPM8 utilizes a semi-swap arrangement as part of its gating pathway (fig. S13C).

## Acknowledgments

We thank Ying Yin and Feng Zhang for preliminary AITC and Cd^2+^ accessibility studies on TRPM8. Cryo-EM data were screened and collected at the Duke University Shared Materials Instrumentation Facility (SMIF), the Pacific Northwest Center for Cryo-EM (PNCC) at OHSU, the National Cancer Institute’s National Cryo-EM Microscopy facility (NCEF) at the Frederick National Laboratory for Cancer Research, and the National Institute of Environmental Health Sciences (NIEHS). We thank Janette Myers at PNCC, Tara Fox at NCEF, and Nilakshee Bhattacharya at SMIF for assistance with the microscope operation. We thank Yang Suo for assistance with data collection at SMIF. This work was supported by the National Institutes of Health (R35NS097241 and R01EY031698 to S.-Y.L.), by the National Institute of Health Intramural Research Program; US National Institutes of Environmental Health Sciences (ZIC ES103326 to M.J.B), and by the NCI’s NCEF under contract 75N91019D00024.

## Funding

National Institutes of Health grant R35NS097241 (S.-Y.L.)

National Institutes of Health grant R01EY031698 (S.-Y.L.)

National Institute of Health Intramural Research Program

US National Institutes of Environmental Health Sciences grant ZIC ES103326 (M.J.B)

NCI’s NCEF under contract 75N91019D00024

## Author contributions

H.-J.L. conducted all biochemical preparation, cryo-EM experiments, and single-particle 3D reconstruction under the guidance of S.-Y.L. C.-G.P. carried out electrophysiological recordings under the guidance of S.-Y.L. J.F. performed data analysis. W.A.P. helped optimize freezing conditions under the guidance of M.J.B. H-J.L. and S.-Y.L. performed model building. S.-Y.L. and J.F. wrote the paper with inputs from H-J.L. and C.G.P.

## Data and materials availability

For PIP₂–menthol–TRPM8 in C_0-1_^M^ state, PIP₂–menthol– TRPM8 in C_1_^M^ state, PIP₂–menthol–AITC–TRPM8 in C_1-2_^M^ state, PIP₂–menthol–TRPM8-Ile^846^Val in C_2_^M^ state, PIP₂–AITC–menthol–TRPM8-Ile^846^Val in O_p_^M^ state, PIP₂–AITC–TRPM8 in C_1_^A^ state, and PIP₂–AITC–menthol–TRPM8-Ile^846^Val in O^M^ state, the coordinates have been deposited in the Protein Data Bank with the accession codes XXXX, XXXX, XXXX, XXXX, XXXX, XXXX, and XXXX. The corresponding cryo-EM maps have been deposited in the Electron Microscopy Data Bank with the IDs EMD-XXXXX, EMD-XXXXX, EMD-XXXXX, EMD-XXXXX, EMD-XXXXX, EMD-XXXXX, and EMD-XXXXX. For PIP₂–menthol–TRPM8 in C_1_^M^ state where rapamycin was not added, only the cryo-EM map is deposited with the ID EMD-XXXXX. Correspondence and requests for materials should be addressed to Seok-Yong Lee.

## Materials and Methods

### Protein expression and purification

The cDNA sequences encoding full-length wild-type (WT) and Ile^846^Val mutant mouse TRPM8 channels were cloned into a modified pEG BacMam vector containing a C-terminal PreScission protease cleavage site, followed by FLAG and 10×His tags ^52^. TRPM8 was expressed in HEK293F suspension cells via baculovirus-mediated transduction. Cells were cultured in Freestyle 293 medium (Life Technologies) and maintained at 37°C with 8% CO₂. Recombinant baculoviruses were generated and amplified following the standard Bac-to-Bac® Baculovirus Expression System protocol (Life Technologies). For expression, 4–6% P3 baculovirus was added to HEK293F cells at a density of less than 3.0 million cells mL⁻¹. After incubation for 18 hours, 10 mM sodium butyrate was added, and the temperature was reduced to 30°C. Cells were harvested after 48 hours.

All purification procedures were conducted at 4°C unless otherwise noted. Harvested cells were resuspended in buffer A (20 mM Tris-HCl pH 8.0, 150 mM NaCl, 5% glycerol, 12 µg mL⁻¹ each leupeptin, pepstatin, and aprotinin, 1.2 mM phenylmethylsulfonyl fluoride, and DNase I) and lysed using a Dounce tissue grinder. Membrane proteins were solubilized by gentle agitation for 1 hour following addition of 1% glycol-diosgenin (GDN; Anatrace). Insoluble debris was removed by centrifugation at 16,000 × g for 30 min. The clarified supernatant was incubated with anti-FLAG M2 resin (Sigma-Aldrich) for 40 min with gentle rotation. Resin-bound proteins were transferred to a gravity-flow column (Bio-Rad) and washed with 10 column volumes (CV) of buffer B (20 mM Tris-HCl pH 8.0, 300 mM NaCl, 0.005% GDN, 10 mM ATP, 10 mM MgCl₂) followed by 10 CV buffer C (20 mM Tris-HCl pH 8.0, 150 mM NaCl, 0.005% GDN). ATP was included to dissociate nonspecifically bound HSP70 proteins. Proteins were eluted using buffer C supplemented with 0.128 mg mL⁻¹ FLAG peptide. Eluted proteins were concentrated and further purified using a Superose 6 Increase column (Cytiva Life Sciences) equilibrated with buffer C. Peak fractions containing TRPM8 WT or Ile^846^Val were pooled, concentrated to approximately 0.4 mg mL⁻¹, and assessed by SDS-PAGE and small-scale cryo-EM screening.

For TRPM8 samples in complex with AITC, allyl isothiocyanate (AITC; Sigma-Aldrich) was incorporated throughout purification. Specifically, harvested cells were resuspended in buffer A, incubated at 37°C for 15 min with gentle rotation, then supplemented with 250 µM AITC and incubated for an additional 30 s at room temperature. Samples were rapidly cooled to below 8°C within 5 min, after which subsequent purification steps were identical to those described above.

### Cryo-EM grid preparation

All grid preparation procedures were performed at 4°C unless otherwise specified. The PIP₂– menthol–TRPM8-Ile^846^Val sample was prepared at 20°C (ambient temperature), while grids were prepared at 4°C and 20°C for the PIP₂–menthol–AITC–TRPM8-Ile^846^Val. For samples vitrified at 20°C, the protein samples were equilibrated at 20°C for at least 10 min prior to ligand incubation.

Freshly concentrated proteins were incubated with 1 mM water-soluble diC8-PIP₂ (Echelon Biosciences) for at least 30 min prior to vitrification. For the PIP₂–AITC–TRPM8 sample (C_1_^A^), protein was incubated with 1 mM diC8-PIP₂ and additional 250 µM AITC. For the PIP₂– menthol–TRPM8 sample (C_1_^M^), protein was incubated with 1 mM diC8-PIP₂ and 1 mM menthol (Sigma-Aldrich). For the PIP₂–menthol–rapamycin–TRPM8 sample (C_0-1_^M^ and C_1_^M^), protein was incubated with 1 mM diC8-PIP₂, 1 mM menthol, and 200 µM rapamycin (Thermo Fisher Scientific). For the PIP₂–menthol–AITC–TRPM8 sample (C_1_^M′^), protein was incubated with 1 mM diC8-PIP₂, 1 mM menthol, and additional 250 µM AITC. For the PIP₂–menthol–TRPM8-Ile^846^Val sample (C_2_^M, 20°C^), protein was incubated with 1 mM diC8-PIP₂ and 1 mM menthol. For the PIP₂– menthol-AITC–TRPM8-Ile^846^Val sample (C_2_^M^, O^M^, and O_p_^M, 20°C^), protein was incubated with 1 mM diC8-PIP₂, 1 mM menthol, and additional 250 µM AITC. All ligands were incubated with protein samples for 3–5 min immediately before grid freezing.

Cryo-EM grids were prepared by applying 3 µL of protein onto freshly glow-discharged Quantifoil R 1.2/1.3, 300-mesh copper holey carbon grids, each with an additional 2 nm continuous carbon support layer (Quantifoil). To minimize preferred orientation, 50 µM fluorinated octyl maltoside (FOM; Anatrace) was added directly to the sample-loaded grids immediately before blotting. Grids were blotted for 4 s at blot force 0 and plunge-frozen into liquid ethane using a Mark IV Vitrobot (Thermo Fisher Scientific). For vitrification at 4°C or 20°C, chamber is maintained with 90% or 80% humidity, respectively. Following vitrification, cryo-EM grids were stored under liquid nitrogen until data acquisition.

### Cryo-EM data acquisition and processing

Cryo-EM data for the PIP₂–menthol–AITC–TRPM8 (C_1_^M^′) and PIP₂–menthol–AITC–TRPM8-Ile^846^Val (C_2_^M^ and O^M^) vitrified at 4°C were acquired using a Titan Krios electron microscope (Thermo Fisher Scientific) operating at 300 keV, equipped with a K3 direct electron detector (Gatan) and a GIF BioQuantum energy filter (20 eV slit width; Gatan), in counting mode. Data were collected using Latitude-S automated data acquisition software (Gatan), as summarized in Table S1. Movies were recorded at a nominal magnification of 105,000×, corresponding to a pixel size of 0.844 or 0.843 Å pixel⁻¹, and a defocus range of –0.7 to –2.2 µm. Movies were not collected in super resolution mode. Each movie stack comprised 40 frames and was exposed to a total electron dose of approximately 60 e⁻ Å⁻².

For other samples, datasets were similarly collected on a Titan Krios microscope equipped as described above, at a magnification of 105,000× with a pixel size of 0.855 Å pixel⁻¹, defocus range of –0.75 to –2.0 µm, and total dose of ∼50 e⁻ Å⁻² per 40-frame movie stack. The dataset for PIP₂–menthol–rapamycin–TRPM8 (C_0-1_^M^ and C_1_^M^) was collected at a pixel size of 0.835 Å pixel⁻¹, with other parameters identical to those described above. Detailed acquisition conditions are summarized in Table S1.

Cryo-EM data processing was carried out as outlined in fig. S3 using RELION 4.0 and CryoSPARC software ^53,54^. Beam-induced motion correction and dose weighting were performed with RELIONs own implementation. Contrast transfer function (CTF) were estimated using the Patch CTF estimation in CryoSPARC, and micrographs were manually curated based on CTF quality and estimated resolution. Particles were initially selected via template-based picking, using 2D templates generated from randomly selected micrographs, followed by iterative rounds of two-dimensional (2D) classification and particle picking refinement using Topaz training. Multiple rounds of 2D classification were conducted to remove false positives, poorly aligned particles, and impurities such as heat shock proteins. Subsequent heterogenous refinement steps utilized a previously published 3D map (EMD-27892) as initial reference, imposing C4 symmetry and optimized parameters (batch size per class 5000, O-EM learning rate initial 1, halflife 100). Final particles used for 3D consensus map reconstruction of TRPM8 showing reasonable feature of side view were transferred to RELION. Usually, densities corresponding to pore domain of TRPM8 are ambiguous, necessitating further classification.

To enhance pore-domain resolution, signals outside the transmembrane domain (TMD) were subtracted, and TMD-focused 3D classification without alignment was performed with varying parameters (K=2–6, T=10 or 20) (fig. S3B). The tight mask for signal subtraction excluded the detergent micelle and cytoplasmic domains (fig. S3B). Particles belonging to high-quality classes or distinct TMD conformations were reverted to original particle and subjected to 3D auto-refinement, followed by CTF refinement and Bayesian polishing in RELION. Final refined particles were imported into CryoSPARC for non-uniform refinement and local refinement to generate the final reconstructions.

Local resolution estimations (fig. S5A), Fourier shell correlation (FSC) validations (fig. S4B,D) based on the gold-standard FSC criterion (0.143 and 0.5), and viewing direction distribution (fig. S4C) were calculated in CryoSPARC. Comprehensive data processing parameters and statistics are summarized in Table S1.

### Model building and refinement

The previously published structures of PIP₂–C3–AITC–mouse TRPM8 (PDB: 8E4L) and PIP₂– mouse TRPM8 (PDB: 8E4N) served as initial models for the mouse Ile^846^Val dataset and all other datasets, respectively. Models were rigid-body fitted into well resolved EM density maps and side chains were manually adjusted to optimal rotamer conformations, and loop regions were rebuilt to accurately fit into densities. Residues with bulky side chains guided the correct registration of helices and β-strands. Special attention was paid to residue registration around the S6 gate region, guided by high-quality, continuous EM density. Throughout manual model building in Coot, ideal geometry restraints were applied to secondary structures and rotamer conformations ^55^. Ligand geometry restraint files were generated from canonical SMILES strings using the eLBOW tool within PHENIX ^56^.

Structural models were validated by calculating Fourier shell correlation (FSC) curves between EM maps and corresponding final atomic models (fig. S4D). Model quality was assessed using MolProbity^57^ within PHENIX, and detailed model refinement and validation statistics are presented in Table S1. Pore radii and surfaces along the channel pathway were calculated using the HOLE program ^58^. Structural analyses, alignments, and illustrations were performed using Coot, UCSF ChimeraX, and PyMOL (Schrödinger) ^59–61^. For comparative analyses presented in Fig. 4, structures were aligned based on VSLD residues 734–853 using PyMOL’s “align” command. Visualization and analysis of cryo-EM reconstructions and EM densities were performed with UCSF Chimera, UCSF ChimeraX, and PyMOL.

### HEK293T cell transfection and whole-cell patch-clamp electrophysiology

HEK293T cells were cultured in DMEM supplemented with 10% fetal bovine serum (Gibco) and 1% Antibiotic-Antimycotic (Gibco) at 37°C in a 5% CO₂ incubator. Cells were plated onto 35 mm dishes and transiently transfected when they reached approximately 50% confluency, with 1 µg of wildtype or mutant constructs of mouse TRPM8, human TRPM2, mouse TRPM3, or human TRPM4 in pBacMam vector and 0.1 µg eGFP, using X-tremeGENE™ DNA transfection reagent (Roche). 1-2 days after the transfection, whole-cell patch clamp recordings were done at room temperature (∼20–22 °C). Cold stimuli were achieved by passing the external recording solution through glass capillary spirals immersed in an ice-water bath maintained at about 0°C, and recordings were performed during constant perfusion with temperature measured using a thermistor (TA-29, Warner Instruments) located close to the cell. The thermistor was connected to the digitizer via a temperature controller (CL-100, Warner Instruments). L-Menthol (Sigma Aldrich) and ruthenium red (RR; Sigma Aldrich), and AMG2850 (Alomone Labs) were applied using a gravity-fed perfusion system for the corresponding measurements. Data were acquired with an Axopatch 200B amplifier (Molecular Devices) and currents were low-pass filtered at 2 kHz (Axopatch 200B) and digitally sampled at 5–10 kHz (Digidata 1440 A). Recording pipettes were pulled from borosilicate glass using a Sutter P-1000 puller and heat-polished to achieve a tip resistance of 2-4 MΩ. 90% series resistance (Rs) compensation was used in all whole-cell recordings. The extracellular solution consisted of 140 mM NaCl, 1 mM MgCl_2_, 10 mM HEPES, and adjusted to pH 7.4 with NaOH. Electrodes were filled with an intracellular solution containing 140 mM CsCl, 1 mM MgCl_2_, 10 mM HEPES, and adjusted to pH 7.4 with CsOH. After breaking the cell membrane, the cell was allowed to equilibrate with the pipette solution for 2–3 min before recording was carried out. All the electrophysiological data analyses were done using Igor Pro 6.2 (Wavemetrics).

### Surface expression biotinylation assay

HEK293T transfections of WT and mutant hTRPM8 were conducted as described above. Transfected cells were cultured in 6-well plates to near confluency. Surface biotinylation was conducted on ice or at 4°C, similarly as previously reported ^62^. Prior to biotinylation, DMEM was aspirated off and the cells carefully rinsed thrice with 1 mL ice-cold DPBS (+Ca^2+^/Mg^2+^). Biotinylation was initiated by replacement with 1 mL DPBS freshly supplemented with 0.5 mg mL^-1^ EZ-link Sulfo-NHS-SS-Biotin (Thermo Scientific), then incubated for 30 min at 4°C. The reaction was then quenched by two 5-minute rinses with 1 mL ice-cold DPBS +100 mM glycine. Cells were lysed by replacing the quench solution with 750 µL of extraction buffer to each well (20 mM Tris-HCl (pH 8.0), 150 mM NaCl, 1%(w/v) GDN, 5 mM EDTA, 10 µg mL^-1^ each of aprotinin, leupeptin and pepstatin, and 0.2 mM PMSF). Cell lysates were disrupted by pipet then transferred to a 1.7 mL tube and left to extract for 1h at 4°C on a rotator. Insoluble material was pelleted at 16,900xg for 15 min at 4°C. The clarified lysates were sampled for protein quantification by bicinconninic acid assay and a consistent, maximal amount of samples were incubated with 50 µL pre-equilibrated High Capacity Neutravidin resin (Pierce) and left to bind for 1 hr at 4°C on a rotator. The resin was then pelleted, washed four times with 1 mL wash buffer (lysis buffer +150 mM NaCl). Afterward, excess wash buffer was aspirated off the beads and the bound samples were eluted with 35 µL 2x loading sample buffer +100 mM DTT then incubated at 65°C for 10 min prior resolution by SDS-PAGE. Semi-dry transfer to 0.45 µm PVDF membranes was followed by blocking with 5% BSA in Tris-buffered saline +0.1% Tween-20 (TBST) and probed by overnight incubation at 4°C with 1,000x diluted monoclonal mouse anti-FLAG M2 antibody (Sigma Aldrich), then 1 hr incubation at room temperature with 10,000x polyclonal goat mouse IgG HRP-conjugated antibody (Rockland). Following both primary and secondary antibody incubations, membranes were washed four times with TBST (each wash lasting 5 min) to remove non-specifically bound proteins. Images were detected using the SuperSignal^TM^ West Pico PLUS reagent kit (Thermo Fisher).

### Statistical analysis

Statistical analyses were performed with Igor Pro 6.2, Excel office 365, and GraphPad Prism 7.0. To analyze the kinetics of current inhibition, the decay phase of inward currents was fitted with a single exponential function using Igor Pro 6.2. The fitting was based on the equation (1).

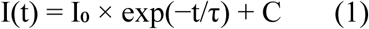

where I(t) is the current at time t, I₀ is the initial amplitude, τ is the time constant representing the rate of inhibition, and C is the steady-state current. All quantitative data are represented as mean ± SEM. Comparisons between two groups were analyzed by Student’s two-tailed unpaired *t*-test. Comparisons among more than two groups were analyzed using one-way ANOVA followed by Dunnett post-hoc test. Comparisons among more than two groups with two independent variables were analyzed using two-way ANOVA followed by Sidak post-hoc test. Differences were considered significant at the *P < 0.05, **P < 0.01, and ***P < 0.001, as appropriate.

## Supplementary Figures

**Fig. S1.**
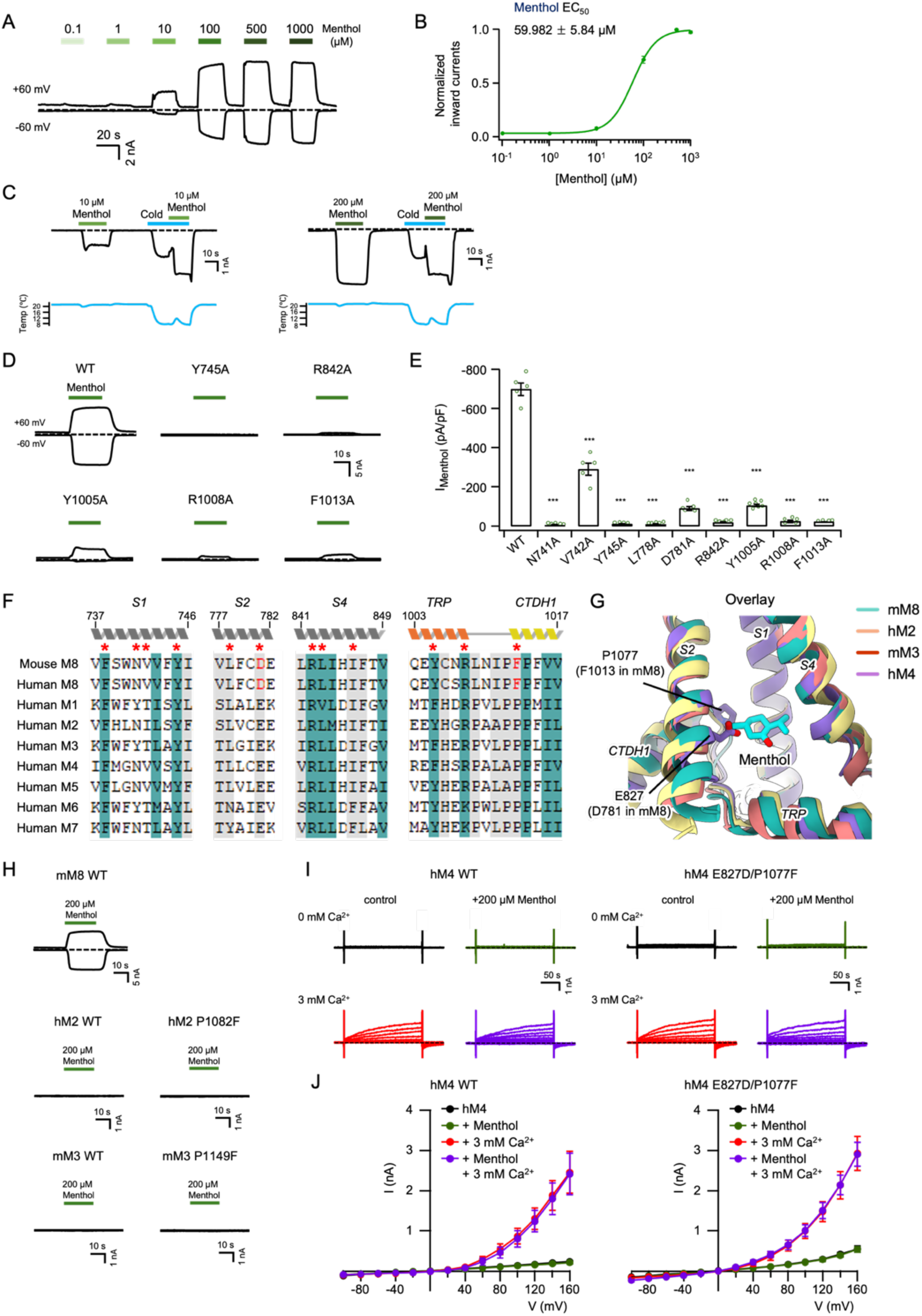
Specificity of menthol binding on TRPM8. **A)** Representative currents of the WT mTRPM8 at -60 mV (lower traces) and +60 mV (upper traces) in HEK293T cells at ∼20°C in response to increasing concentrations of menthol. Dotted lines indicate the zero-current level. **B)** Concentration-response curve of menthol activation measured from whole-cell recordings at - 60 mV (as in **A**), normalized to the maximal response, shows mean ± SEM (*n* = 7 biological replicates). The continuous curves were fit to the Hill equation with EC_50_ values indicated in the figure. **C)** Representative current traces of the WT mTRPM8, elicited by menthol alone or by cold followed by 10 μM or 200 μM menthol at −60 mV in HEK293T cells. Horizontal colored lines represent the application of cold (sky blue), 10 μM menthol (light green), and 200 μM menthol (green), as indicated. Dotted lines indicate the zero-current level. **D)** Representative currents of the mTRPM8 WT and menthol binding site mutants at -60 mV (lower traces) and +60 mV (upper traces) in HEK293T cells. Horizontal colored lines represent the application of 200 μM menthol (green) as indicated. Dotted lines indicate the zero-current level. **E)** Summary of menthol-evoked inward current densities (pA/pF) for mTRPM8 WT and menthol binding site mutants (*n* = 4-7 biological replicates). Cells were transfected with the same amount of cDNA. Dots indicate the biological replicates for each experiment. \*\*\**P* < 0.001, using one-way ANOVA followed by Dunnett’s post-hoc test. Data are mean ± SEM. **F)** Sequence alignment of human TRPM channels. Residues with 100% and 80% conservation are shaded in green and light grey, respectively. Residues involved in menthol binding within the VSLD are marked with red asterisks. Asp^781^ and Phe^1013^, highlighted in red, are uniquely conserved in TRPM8 among the TRPM family. **G)** Structural alignment of mTRPM8 (C_1_^M’^) menthol binding site against human TRPM2 (PDB 6PUO)^63^, mouse TRPM3 (PDB 9B2A)^64^ and human TRPM4 (PDB 6BQV)^65^, as indicated. Only Phe^1013^ and Asp^781^ differ in sequence (Fig. S1F), being Pro^1077^ and Glu^827^ in hTRPM4, respectively. **H)** Representative currents of the WT and mutant forms of hTRPM2 and mTRPM3 at -60 mV (lower traces) and +60 mV (upper traces) in HEK293T cells. The mTRPM8 current trace (previously shown in fig. S1D) is included here as a reference for comparison. Horizontal colored lines represent the application of 200 μM menthol (green) as indicated. Dotted lines indicate the zero-current level. **I)** Currents elicited between -100 mV and +160 mV in 20 mV steps in HEK293T cells expressing hTRPM4 WT and Glu^827^Asp/Pro^1077^Phe double-mutant. Holding potential is -60 mV. Representative current traces obtained in the absence or presence of 200 μM menthol and 3 mM Ca^2+^, with conditions color-coded as follows: black traces for control (absence of menthol and Ca^2+^), green traces for menthol alone, red traces for Ca^2+^ alone, and purple traces for both menthol and Ca^2+^. **J)** Currents elicited between -100 mV and +160 mV in 20 mV steps in HEK293T cells expressing hTRPM4 WT and Glu^827^Asp/Pro^1077^Phe double-mutant. Holding potential is -60 mV. Current-voltage (I-V) curves of WT and Glu^827^Asp/Pro^1077^Phe double-mutant hTRPM4 (*n* = 4-5 biological replicates) obtained in the absence or presence of 200 μM menthol and 3 mM Ca^2+^, with conditions color-coded as follows: black for hM4 (absence of menthol and Ca^2+^), green for menthol alone, red for Ca^2+^ alone, and purple for both menthol and Ca^2+^. Data are mean ± SEM.

**Fig. S2.**
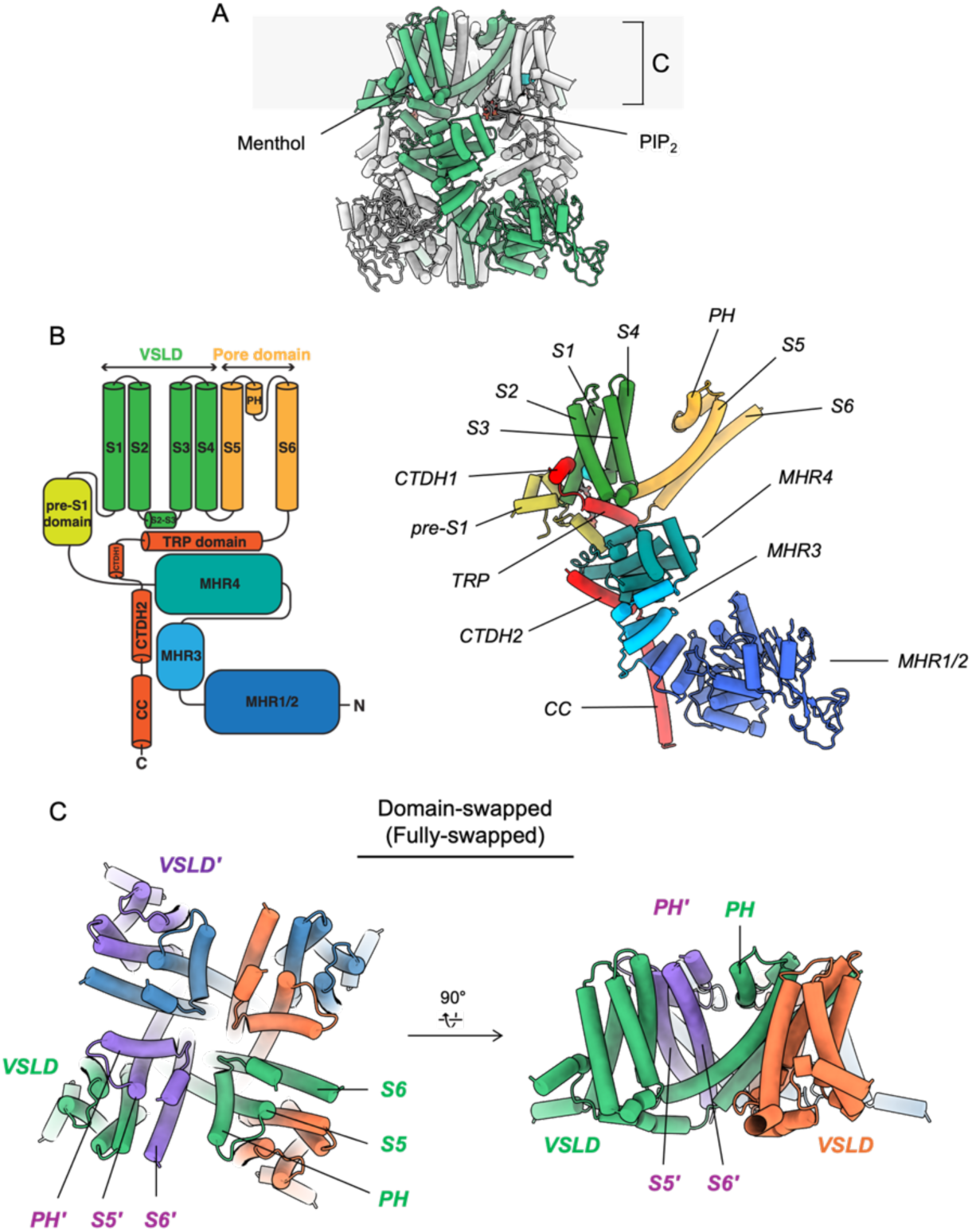
Overall architecture of TRPM8. **A)** Cartoon representation of the TRPM8 tetramer in a domain-swapped conformation. **B)** Protomer topology illustrated schematically (left) and structurally represented with cylindrical helices (right). Structural representation (right) is colored consistently with the schematic (left). **C) Structure of TMD region** is depicted as cartoons, with neighboring protomers distinctly colored to highlight that the pore domain from one protomer is intercalated between the VSLD and pore domain of its neighbor (fully-swapped).

**Fig. S3.**
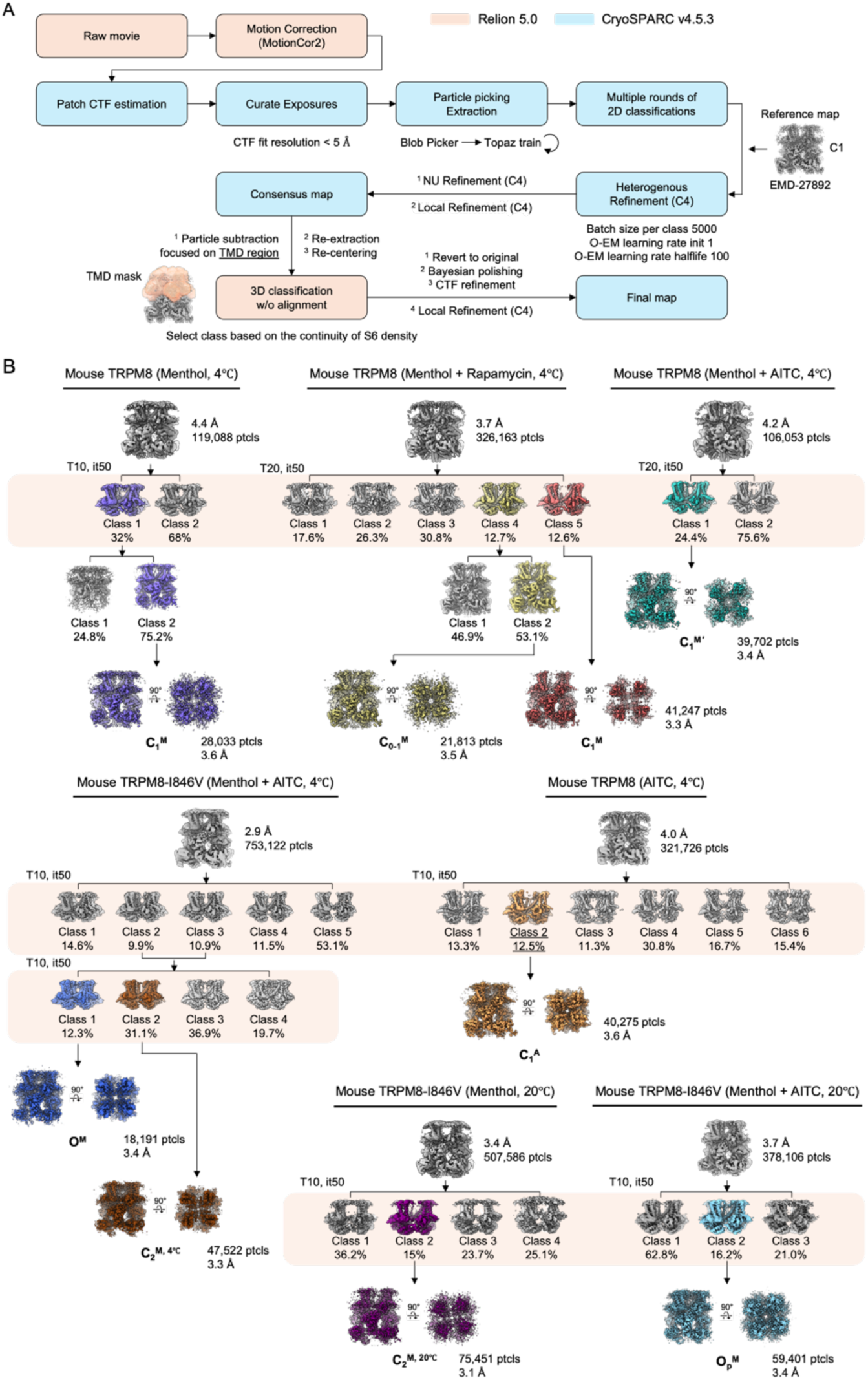
Cryo-EM data processing workflow. **A)** Overview of the cryo-EM image processing pipeline employed to determine TRPM8 structures presented in this study. Due to the intrinsic flexibility of the pore domain, the workflow specifically targets heterogeneity within the transmembrane domain (TMD) through focused 3D classification without alignment. Classes for subsequent processing were selected based on the clear continuity of the S6 helix density. **B)** For all datasets, TMD-focused classification was performed using varying parameters in RELION, followed by final map reconstruction in CryoSPARC. Detailed procedures are described in the Methods.

**Fig. S4.**
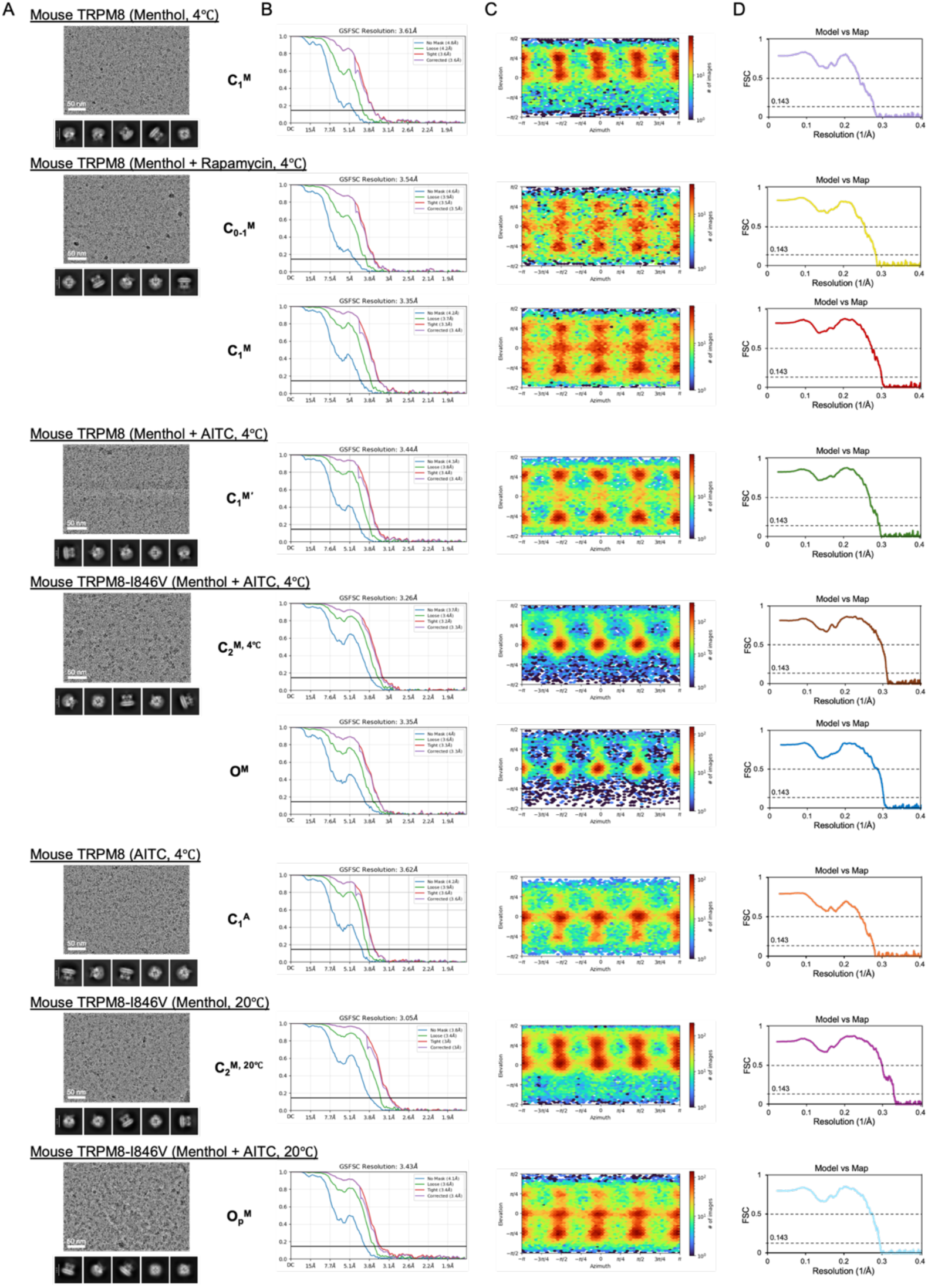
Representative micrographs, 2D class averages, and FSC curves. **A)** For each dataset, a representative micrograph, five 2D class averages, a plot showing the distribution of particle orientations, and a Fourier shell correlation (FSC) plot of the final map are presented from left to right, respectively. FSC curves between two half-maps were calculated using the 0.143 criterion. **B)** For each dataset, FSC curves comparing the final model to the corresponding map are shown.

**Fig. S5.**
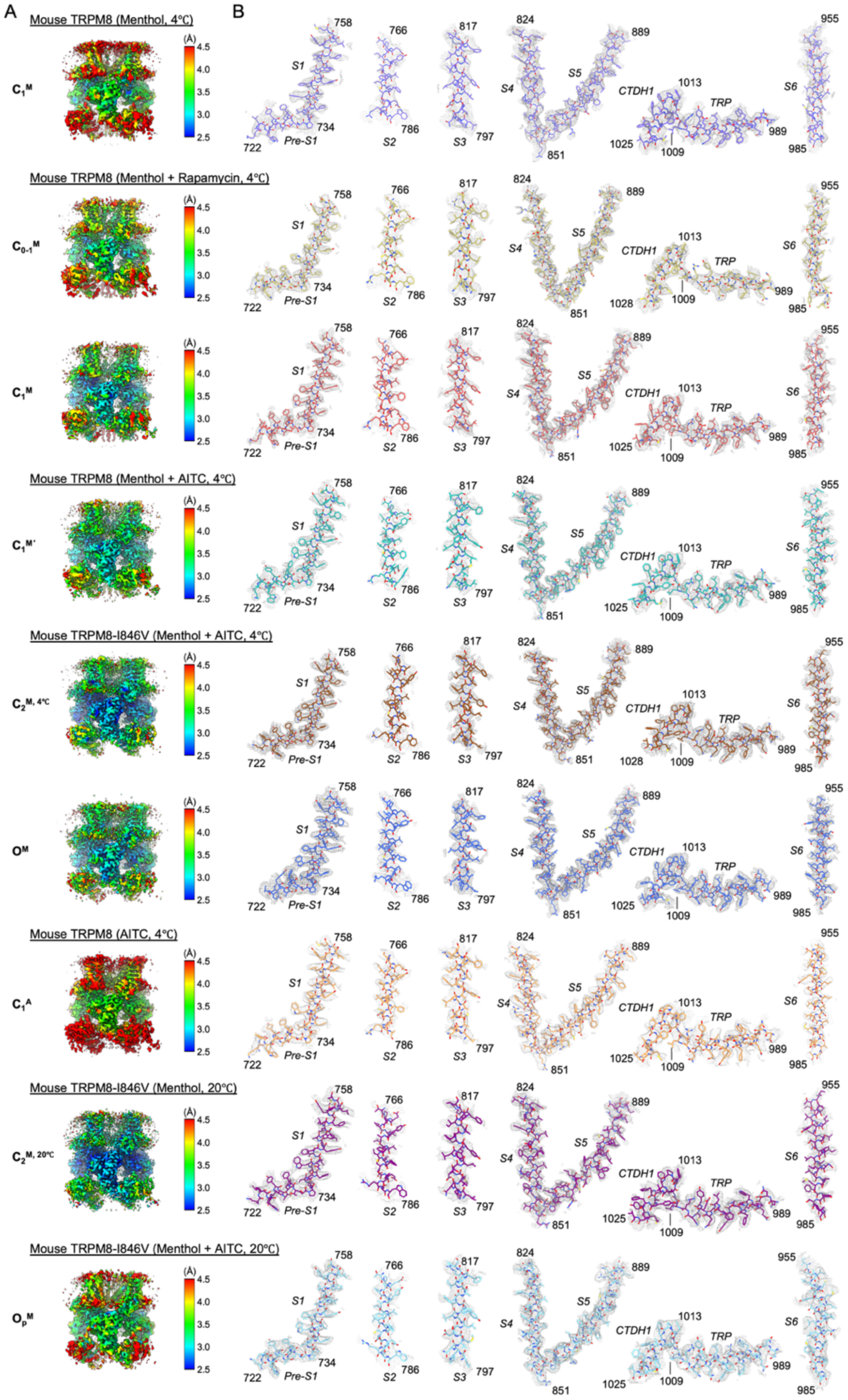
Quality of cryo-EM maps. **A)** Final cryo-EM maps colored according to local resolution estimates. The color scale indicates local resolution in Å, as shown in the accompanying bar. **B)** Densities corresponding to the transmembrane domain (TMD) regions of each structure are displayed as grey meshes overlaid with the corresponding atomic models.

**Fig. S6.**
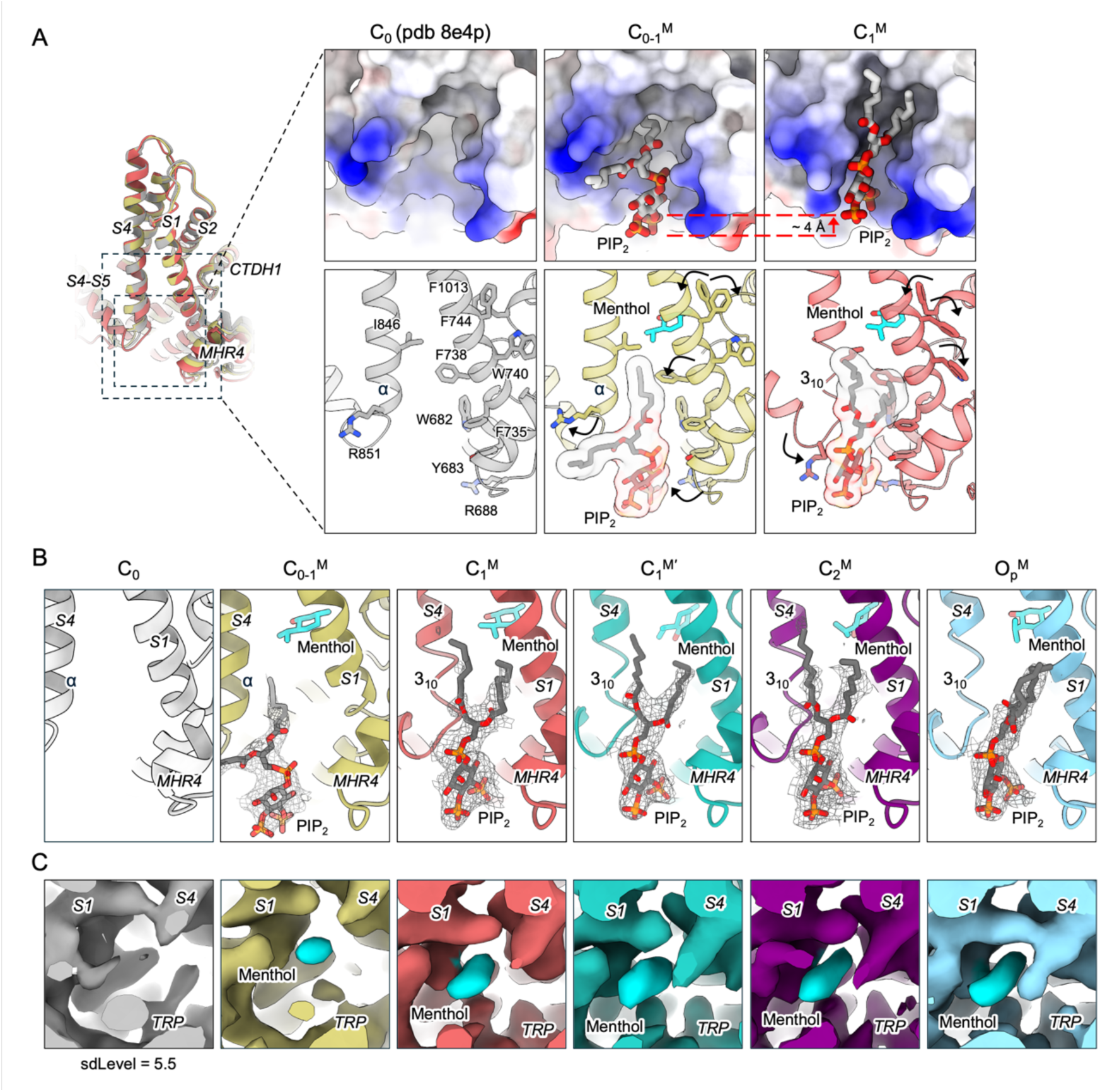
PIP₂ and menthol binding modes across different states. **A)** Overlay of TRPM8 protomer structures in the C_0_ (PDB 8E4P; grey), C_0-1_^M^ (yellow), and C_1_^M^ (red) states, shown as cartoons (left). Surface representations highlight the electrostatic properties of each state and illustrate distinct PIP₂ (grey sticks) binding modes within the interfacial cavity (top right). Upon transition from C_0-1_^M^ to C_1_^M^, PIP₂ shifts ∼4 Å toward the membrane and engages more tightly. The lower right panel presents PIP₂ with its corresponding transparent surface from the same viewpoint. Residues undergoing structural rearrangements are depicted as sticks and labeled. The cytoplasmic half of S4 (S4b) is shown as an α-helix in C_0_ and C_0-1_^M^, and as a 3₁₀-helix in C_1_^M^. Menthol is shown as cyan sticks. One-sided arrows indicate side-chain rotations. **B)** EM densities corresponding to PIP₂ from each structure are depicted as grey mesh. Cartoons of C_1_^M^*′* (green), C_2_^M^ (purple), and O_p_^M^ (sky blue) are shown. In these states, S4b adopts a 3₁₀-helical conformation. **C)** EM densities corresponding to menthol from each structure are shown, colored to match the corresponding cartoon representations. All maps were sharpened using the same B-factor (−80 Å ²) and low-pass filtered to 3.5 Å.

**Fig. S7.**
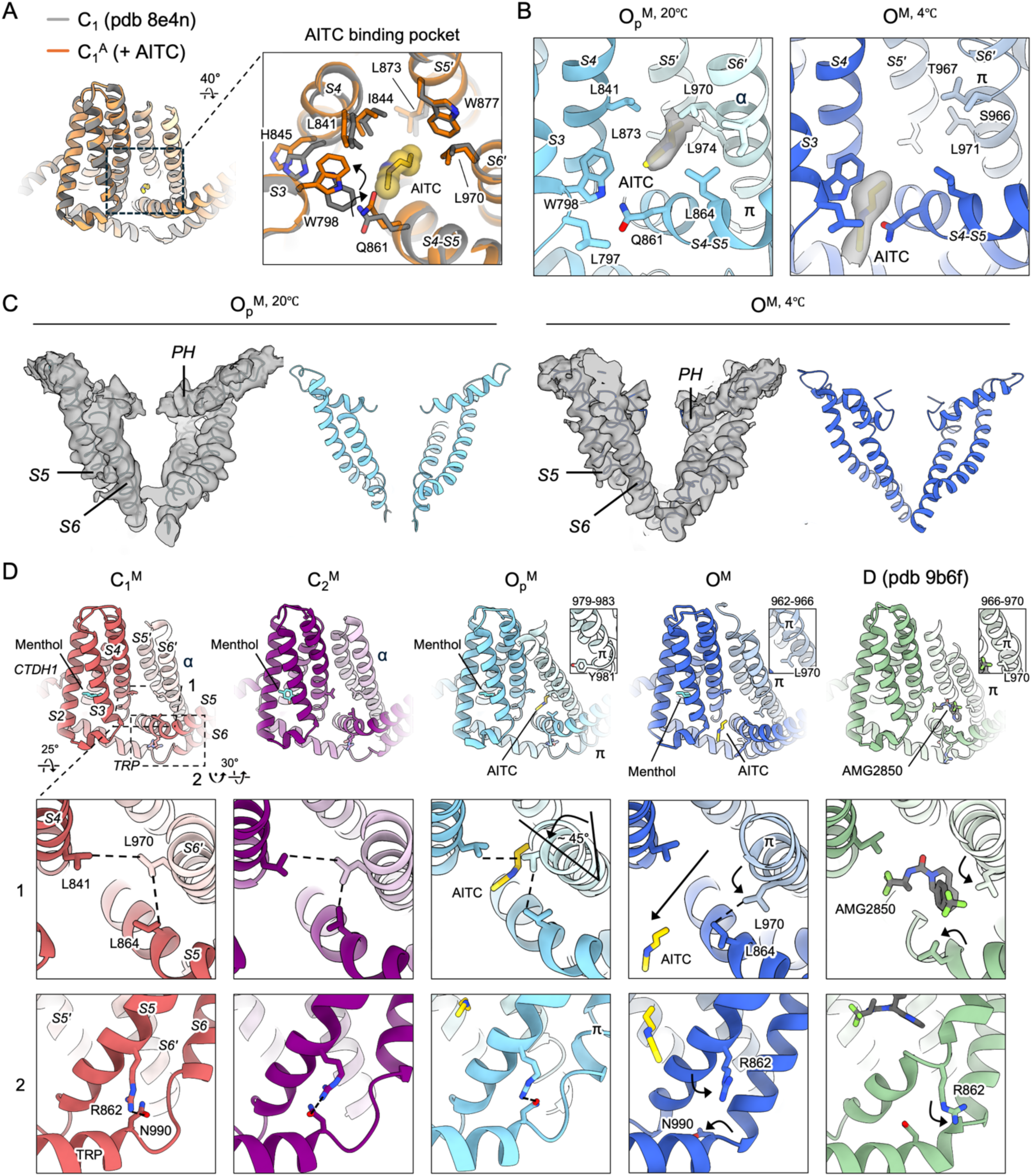
AITC binding pocket and pore helix of O_p_^M^ and O^M^, and changes in interactions at the neighboring interface among different states. **A)** Overlay of neighboring protomers in the C_1_^A^ (orange) and C_1_ (PDB 8E4N; grey) states, shown as cartoons (left). Structural comparison of the AITC binding pocket between C_1_^A^ versus apo C_1_ shows similar Cα trace with slight conformational changes in sidechain. **B)** Detailed interactions of AITC with surrounding residues in O_p_^M, 20℃^ (sky blue) and O^M, 4℃^ (blue). EM density corresponding to AITC is shown as grey transparent surfaces. **C)** Cα traces of S5-PH-S6 in O_p_^M, 20℃^ and O^M, 4℃^ are shown as cartoons overlayed with transparent EM density, highlighting the notable tilt of PH. Due to poor resolution in this area EM densities shown are from unsharpened maps. **D)** For structural comparison, two neighboring protomers are shown as cartoons for the C_1_^M^ (red), C_2_^M^ (purple), Op^M^ (sky blue), O^M^ (blue), and D (PDB: 9B6F; green) states, with corresponding ligands indicated. Inset boxes (top right) highlight the formation of a π-helix at distinct positions in S6 among the O_p_^M^, O^M^, and D (PDB: 9B6F) states. In view 1, Leu^841^ in S4, Leu^864^ in S4–S5, and Leu^970^ in the neighboring S6 (S6′) are depicted as sticks. Black dashed lines indicate hydrophobic interaction networks, while black arrows highlight rotations of S6 helix and rotation of residues. In view 2, Arg^862^ in S4–S5 and Asn^990^ in the TRP domain are shown as sticks, with putative hydrogen bonds indicated by black dashed lines. This interaction is lost as the channel transitions towards opening and desensitization.

**Fig. S8.**
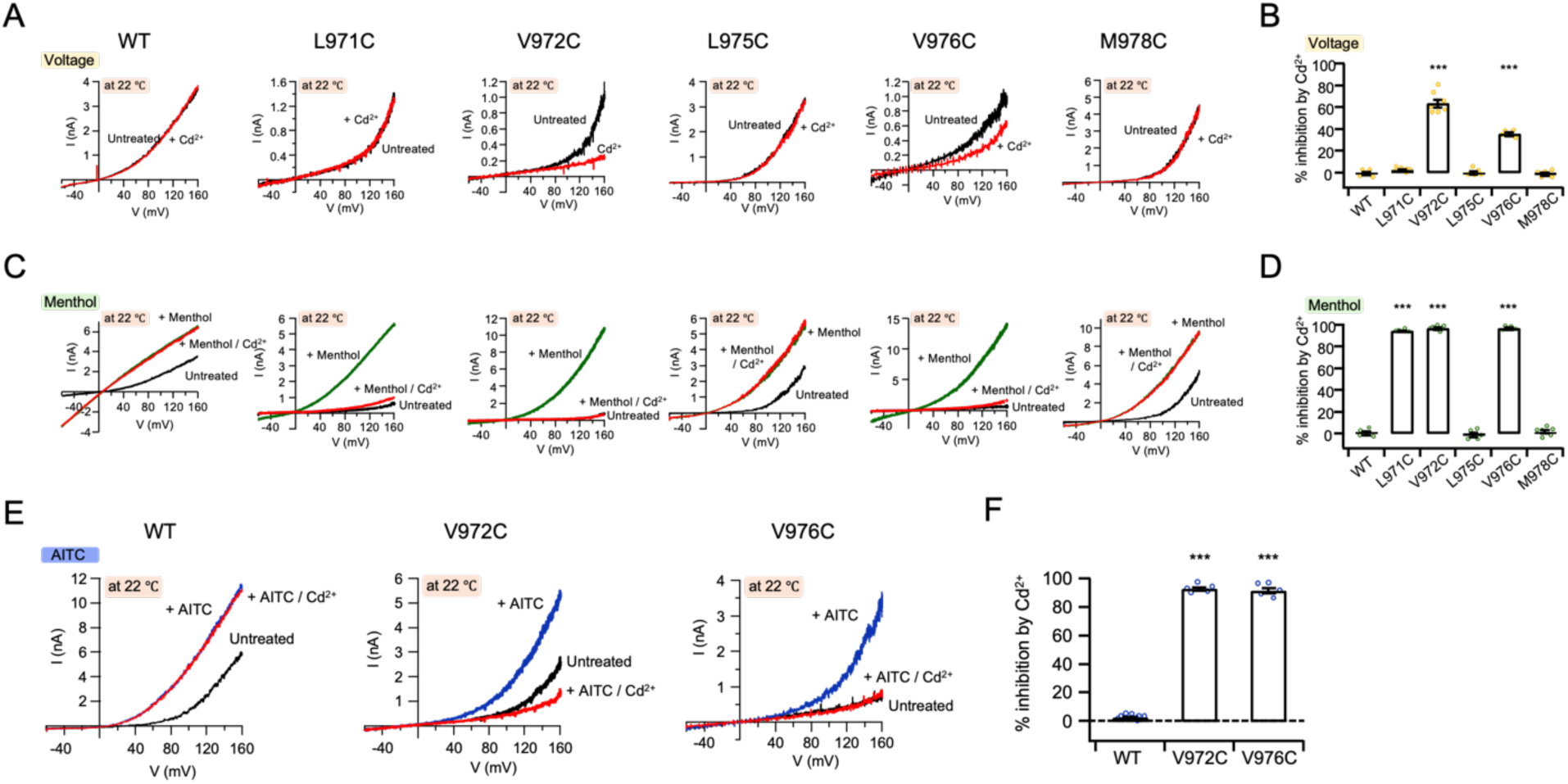
Cd^2+^ block of cysteine mutant scanning of S6. **A)** Representative current-voltage (I-V) plots of the mTRPM8 WT and cysteine mutants in HEK293T cells, obtained using a 300-ms voltage ramp from -60 mV to +160 mV, showing basal currents before any treatment (black trace), followed by 20 μM Cd^2+^(red trace). **B)** Summary of inhibition of basal current by 20 μM Cd^2+^ measured at +160 mV in HEK293T cell with mTRPM8 WT and cysteine mutants (*n* = 4-7 biological replicates). Dots indicate the individual data points for each experiment. \*\*\**P* < 0.001, using one-way ANOVA followed by Dunnett’s post-hoc test. Mean ± SEM are plotted. **C)** Representative current-voltage (I-V) plots of the mTRPM8 WT and cysteine mutants in HEK293T cells, obtained using a 300-ms voltage ramp from -60 mV to +160 mV, showing basal currents before any treatment (black trace), followed by initial application of 200 μM menthol alone (green trace), then co-application of 200 μM menthol with 20 μM Cd^2+^(red trace). **D)** Summary of inhibition of menthol-evoked current by 20 μM Cd^2+^ measured at +160 mV in HEK293T cell with mTRPM8 WT and cysteine mutants (*n* = 4-6 biological replicates). Dots indicate the individual data points for each experiment. \*\*\**P* < 0.001, using one-way ANOVA followed by Dunnett’s post-hoc test. Mean ± SEM are plotted. **E)** Representative current-voltage (I-V) plots of the mTRPM8 WT and cysteine mutants in HEK293T cells, obtained using a 300-ms voltage ramp from -60 mV to +160 mV, showing basal currents before any treatment (black trace), followed by initial application of 2 mM AITC alone (deep blue trace), then co-application of 2 mM AITC with 20 μM Cd^2+^(red trace). **F)** Summary of inhibition of 2mM AITC-evoked current by 20 μM Cd^2+^ measured at +160mV in HEK293T cell with mTRPM8 WT and cysteine mutants (*n* = 5-8 biological replicates). Dots indicate the individual data points for each experiment. \*\*\**P* < 0.001, using one-way ANOVA followed by Dunnett’s post-hoc test.

**Fig. S9.**
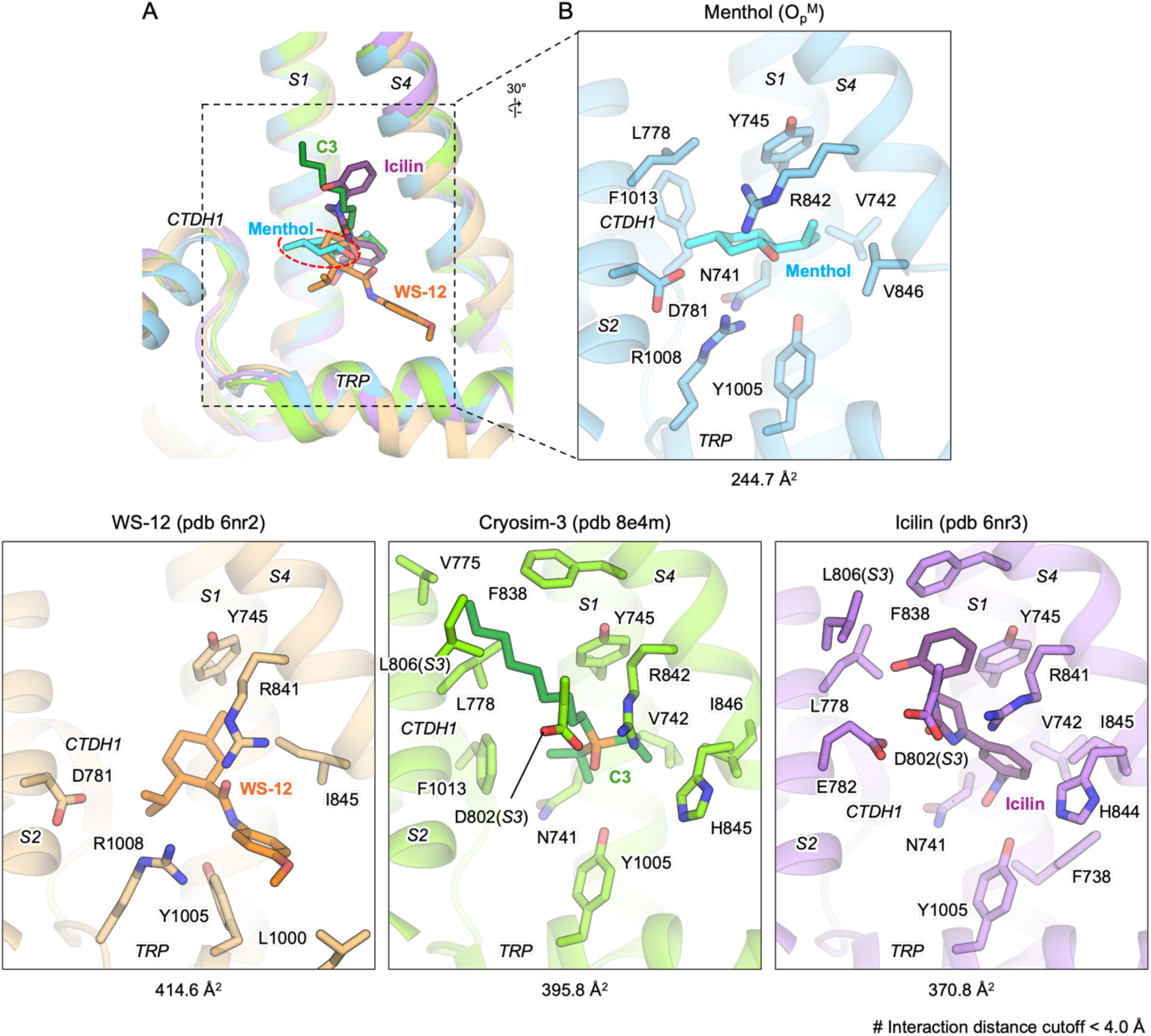
Different cooling agonists bound within the VSLD cavity of TRPM8. **A)** Comparison of binding modes for different cooling agonists within the VSLD cavity. The VSLD regions of menthol (pre-OM; sky blue), WS-12 (PDB 6NR2; orange), Cryosim-3 (PDB 8E4M; green), and icilin (PDB 6NR3; purple) bound structures are shown as cartoons and overlaid. S2, S2–S3, and S3 segments are omitted for clarity. The red dashed circle indicates that menthol is positioned deeper around CTDH1 compared to other agonists. **B)** Residues interacting with each agonist within ∼4 Å are shown as sticks and labeled. The interaction surface area between each agonist and the surrounding protein is indicated below.

**Fig. S10.**
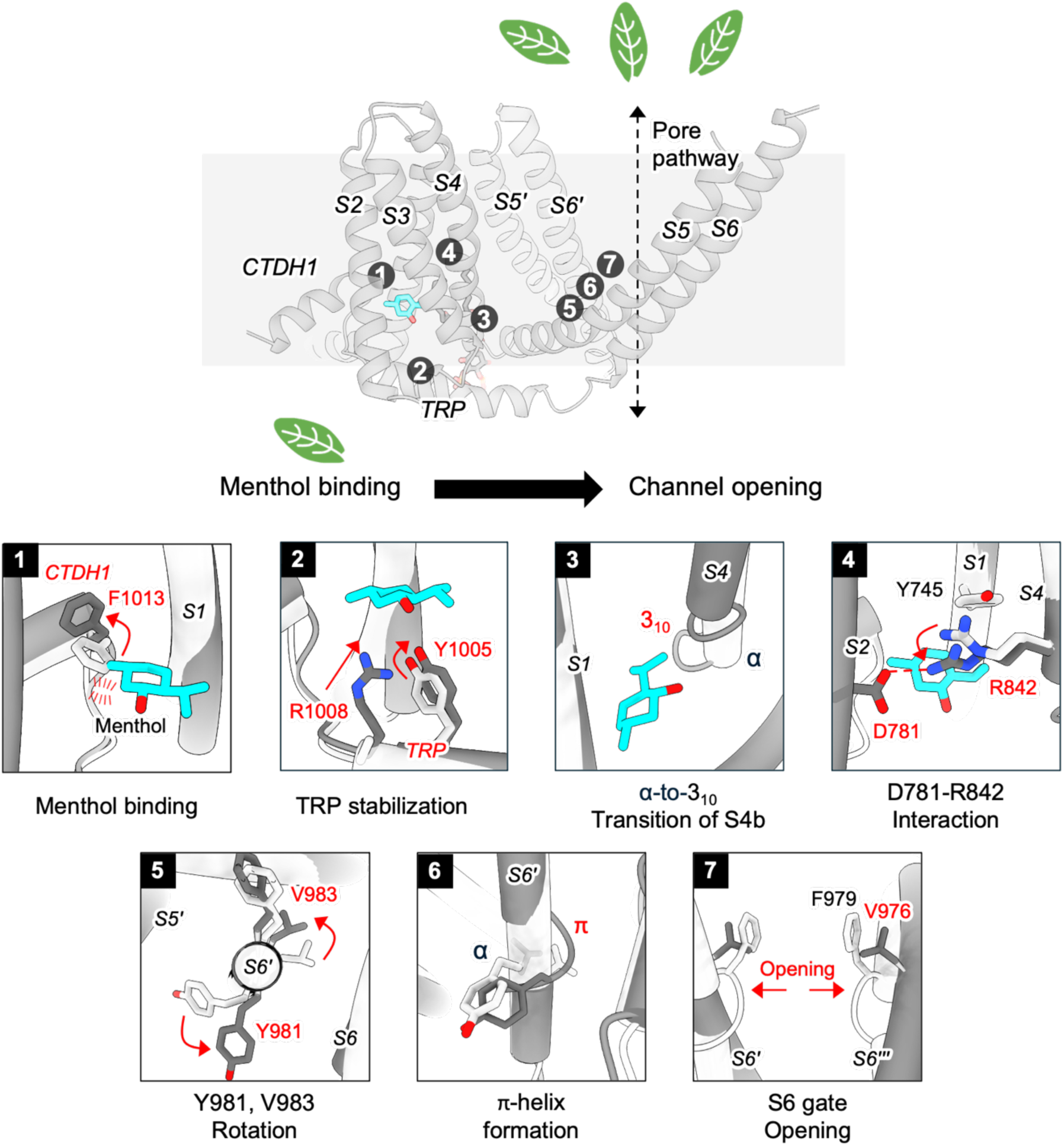
Mechanistic model for menthol and cold induced TRPM8 activation. Serial conformational changes within VSLD induced by menthol binding at 4°C propagates towards S6 gate and opening with interaction changes at the interface between neighboring protomers

**Fig. S11.**
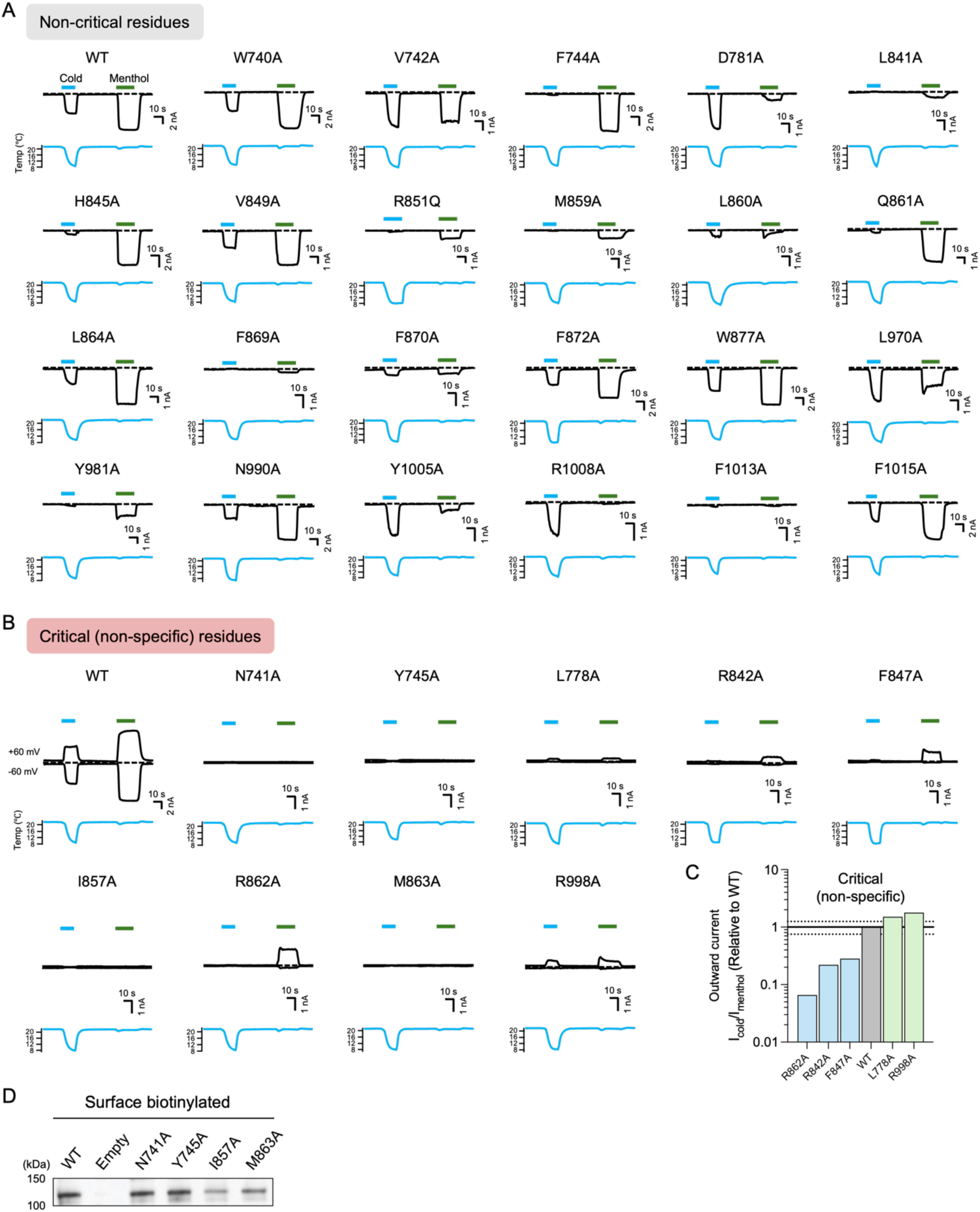
Extensive electrophysiological analysis to identify residues involved in menthol and cold activation. **A)** Representative inward current traces of mTRPM8 WT and alanine-substituted mutants, elicited by cold and 200 μM menthol at −60 mV in HEK293T cells. Horizontal colored lines represent the application of cold (sky blue) and 200 μM menthol (green), as indicated. Dashed lines indicate the zero-current level. **B)** Representative currents of mTRPM8 WT and alanine-substituted “critical (non-specific)” mutants lacking inward-current activity to both menthol and cold. Recordings in HEK293T cells at -60 mV (lower traces) and +60 mV (upper traces) in response to cold (sky blue) and 200 μM menthol (green), as indicated. **C)** The outward current density ratio was used to further categorize five “critical (non-specific)” mutants as menthol- or cold-preferred. **D)** Western blot analysis of surface-biotinylated HEK293T cells transfected with WT TRPM8 and completely non-functional mutants Asn^741^Ala, Tyr^745^Ala, Ile^857^Ala, and Met^863^Ala, demonstrating their proper trafficking to the plasma membrane.

**Fig. S12.**
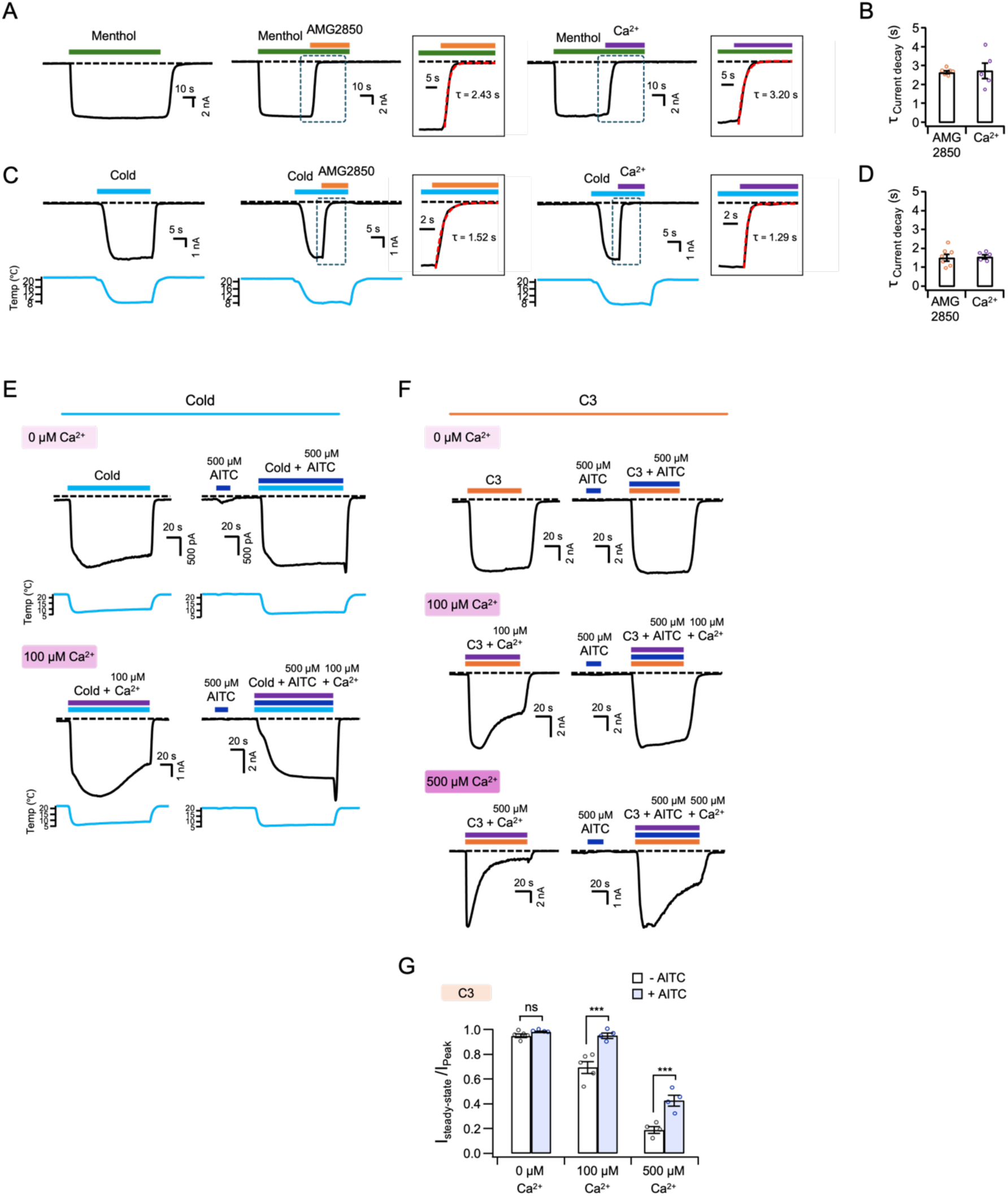
Role of AITC in reducing desensitization of cooling agonist- and cold-activated TRPM8. **A and C)** Representative currents of the WT mTRPM8 channels at –60 mV in HEK293T cells. Current traces elicited by 200 μM menthol (**A**, green) and cold (**C**, sky blue) are shown on the left and are inhibited by subsequent application of 1 μM AMG2850 (middle, orange) and 500 μM Ca^2+^ (right, purple). The segments of current traces used to analyze current decay are highlighted with blue dashed boxes and shown in expanded views on the right, outlined in black. Red dashed curves indicate single exponential fits used to determine the time constants (τ) of current decay. Dotted lines indicate the zero-current level. **B and D)** Summary of time constants (τ) for inhibition of menthol (**B**) and cold (**D**)–activated currents by AMG2850 and Ca²⁺, based on the exponential fitting in **A** and **C**. Dots indicate the individual data points for each experiment with mean ± SEM. **E)** Representative currents of the WT mTRPM8 channels at –60 mV in HEK293T cells. Current traces elicited by cold without (left) or with 500 µM AITC (right) for 2 min in the presence of 0 μM (top) and 100 μM (bottom) extracellular Ca^2+^. **F)** Representative currents of the WT mTRPM8 channels at –60 mV in HEK293T cells. Current traces elicited by 100 μM C3 without (left) or with 500 µM AITC (right) for 2 min in the presence of 0 μM (top), 100 μM (middle), or 500 μM (bottom) extracellular Ca^2+^. **G)** Summary of remaining current after desensitization in the presence of 0 μM, 100 μM, and 500 μM extracellular Ca^2+^ (*n* = 4–5). Dots indicate the individual data points for each experiment with mean ± SEM. ns > 0.05, \*\*\**P* < 0.001, using two-way ANOVA followed by Sidak post-hoc test. Dotted lines indicate the zero-current level.

**Fig. S13.**
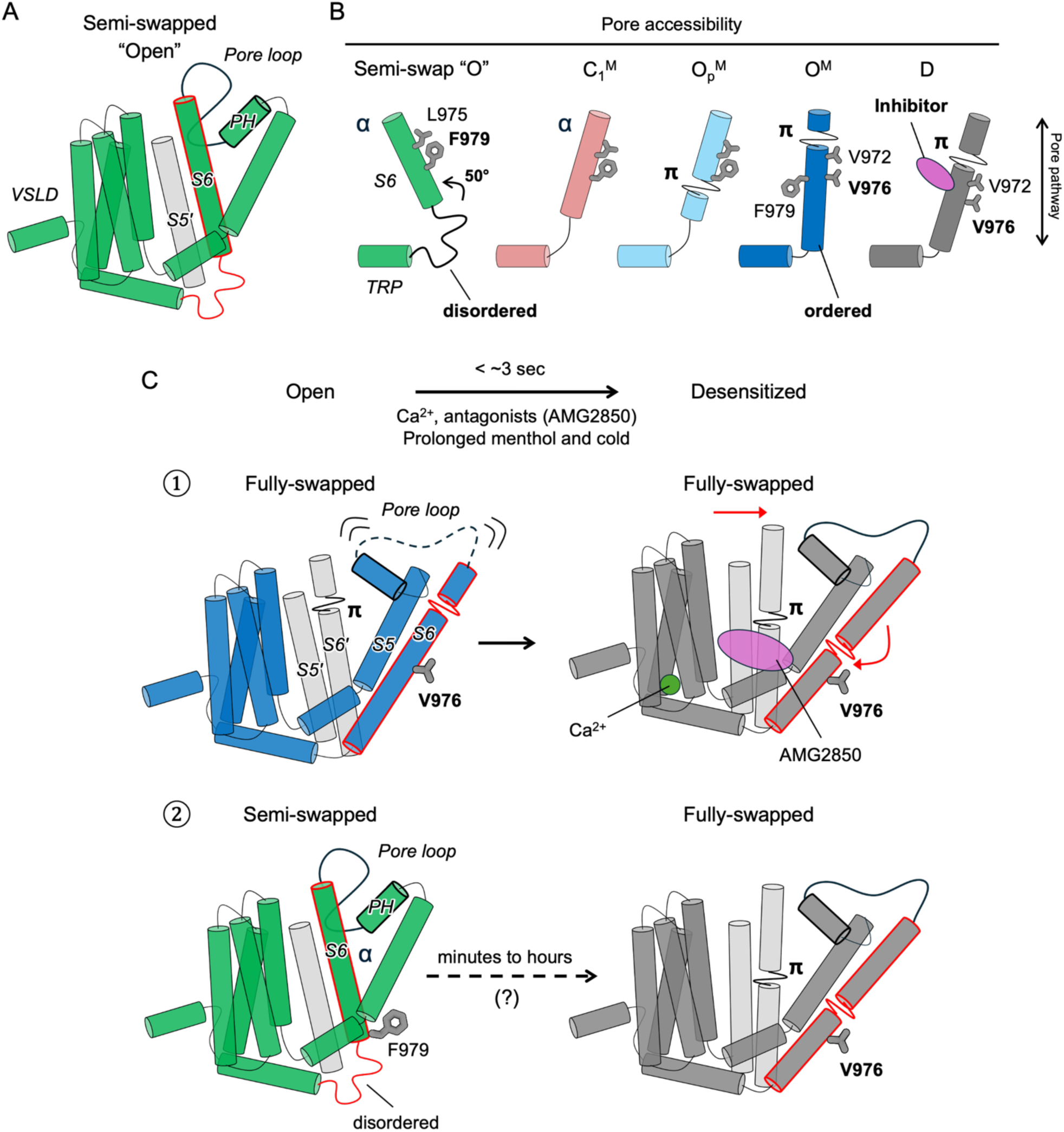
Incompatibility of a semi-swap conformation in TRPM8 gating. **A)** Schematic cartoon of the semi-swapped conformation, recently proposed as an “open” conformation based on figures in the manuscript by Choi. K.Y, *et al.* (2025)^27^. One protomer TMD is shown in green and the S5’ of the adjacent protomer is shown in gray. Red and black outlines respectively highlight S6 and PH positions adjacent to the VSLD of the same protomer and the stabilized pore loop. **B)** Schematic comparison of S6 between semi- and fully-swapped architecture. Relative to the full-swap conformation, S6 in the semi-swap conformation is completely α-helical and is dramatically shorter and tilted by 50° towards the VSLD of same protomer. Consequently, the linker between S6 and TRP helix becomes substantially disordered. In this conformation, L975 and the gate residue F979 are pore-lining and thus surface accessible. On the contrary, in the full-swap conformation, transition from C_1_ to O then D is accompanied by increasing S6 helical propensity toward the TRP helix, rotation and tilting of S6, and formation of the functionally important π-bulge. In the full-swap open conformation, F979 and L975 are functionally verified to be buried while V976 becomes the pore-lining gate. The S6 π-bulge is also important for binding conformationally selective D-state inhibitors, wherein TRPM8 adopts the full-swap conformation. **C)** TRPM8 desensitization towards a single, functionally and structurally verified full-swap conformation is induced by either Ca²⁺ from prolonged menthol or cold activation, or antagonists such as AMG2850. During desensitization, the critical interface comprising S4, the S4-S5 linker, S5’, S6’ and the TRP helix undergoes structural rearrangement with the π-bulge in S6 stabilizing AMG2850 binding. TRPM8 desensitization occurs within 3s (top), consistent with the occurrence of rather small conformational changes. However, the large structural rearrangements in quaternary structure required for transition between semi-swap to full-swap conformations would necessitate partial disassembly of the TRPM8 tetramer in order to accommodate the drastic rotation of S6, PH, and pore loop, which has been measured to occur over several minutes to hours (bottom)^27^. Therefore, activation then desensitization through a semi-swap conformation is incompatible with the functional data, which supports a full-swap architecture of TRPM8.

**Table S1.**
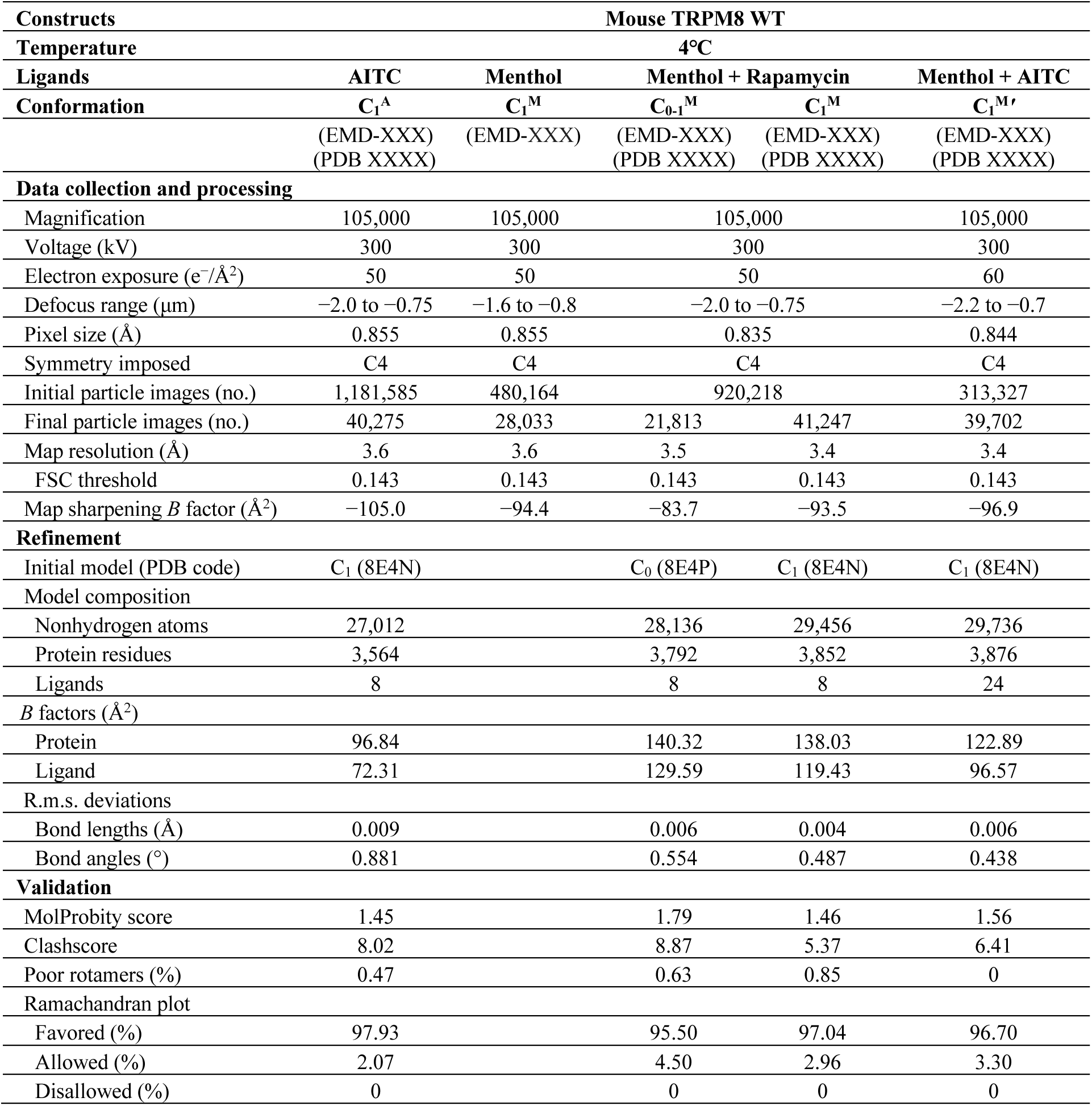
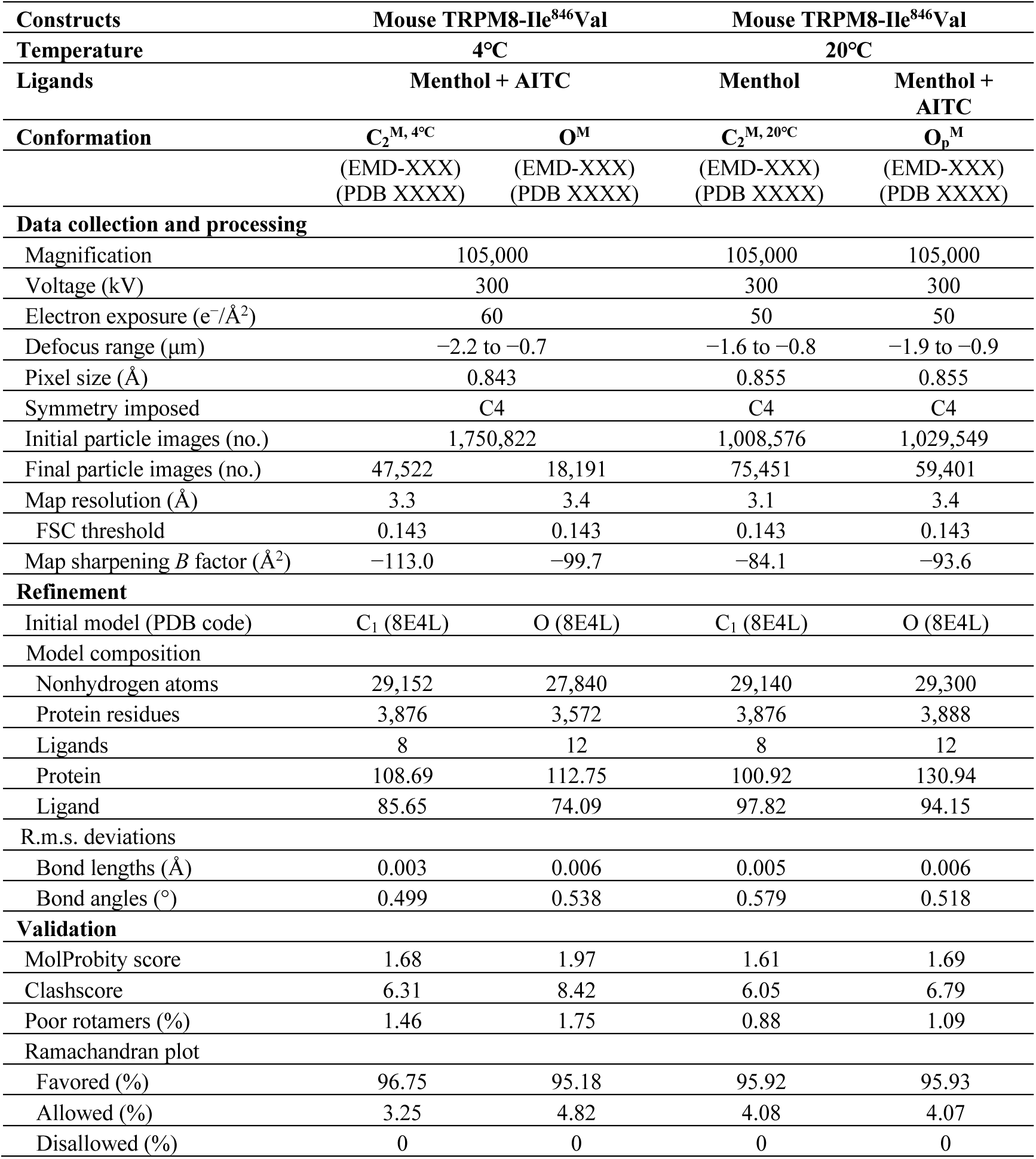
Cryo-EM data collection, refinement, and validation statistics.

